# U1 snRNP-Specific U1C Acts as the Gatekeeper of the Survival of Motor Neurons (SMN) Complex in snRNP Biogenesis

**DOI:** 10.1101/2024.11.11.623077

**Authors:** Duc Minh Ngu, Sanat Myti, Ayesha Ali Khan, Jeannine Keita, Tessa Moore, Paul Andega, Alaa Aziz, Ritu Raj, Meeyeon Park, Kayunta Johnson-Winters, Eul hyun Suh, Jeongsik Yong, Byung Ran So

## Abstract

The stability and abundance of spliceosomal snRNPs are determined by the assembly of an Sm protein ring (Sm core) on each snRNA, a process facilitated by the survival of motor neuron (SMN) complex. Whereas the SMN complex’s role as a chaperone is well-established, the mechanisms that regulate its activity remain unclear. Our study identifies U1C, a U1 snRNP-specific protein, as a key regulator of the SMN complex. U1C together with U1-70K facilitates U1 snRNP formation, rendering the SMN complex accessible for other snRNAs to assemble Sm cores. Remarkably, a prevalent cancer-associated mutation in U1 snRNA near the U1C binding site not only disrupts Sm core assembly but also sequesters the SMN complex, inhibiting canonical snRNP formation. Our findings provide mechanistic insights into how snRNP-specific proteins regulate the SMN complex and suggest that U1 snRNA mutations in numerous cancers disrupt the SMN complex, leading to perturbation in various RNA metabolism pathways.

**Highlights:**

- U1C facilitates the formation of Sm core not only on U1 snRNA but also on other snRNAs.
- U1C together with U1-70K promotes U1 snRNP formation, enabling the SMN complex to recruit Gemin5/other snRNAs complex.
- The divalent binding of U1C to the SMN complex is mediated by U1C’s N-terminus helix, interacting with U1-70K, and its arginine-methylated C-terminus domain, binding to the SMN protein.
- Multiple U1 snRNA mutations in cancer patients defect in Sm core assembly and potentially sequester the SMN complex, thereby inhibiting canonical snRNP formation.

## INTRODUCTION

In most eukaryotes, precursor messenger RNAs (pre-mRNAs) transcribed by RNA polymerase II (Pol II) undergo several processing steps to produce mRNAs. Nascent pre-mRNAs are co-transcriptionally organized by various RNA-binding proteins and non-coding RNAs^1, 2^. One of these intricate steps is splicing, which involves removing introns and ligation of its flanking exons to produce mature mRNAs. This process requires association, rearrangement, and dissociation of multiple components of spliceosomes; mainly small nuclear ribonucleoproteins (snRNPs) and splicing factors^3^. In higher organisms, including humans, alternative splicing is a mechanism that produces a variety of transcript isoforms from a single gene, fine-tuning protein diversity^4, 5^.

Extensive biochemical and structural studies have made significant advances in our understanding of spliceosome composition, structure, and stepwise assembly at the atomic level^6-8^. Although the equal stoichiometry of snRNPs required for splicing is widely accepted, recent studies have shown notable variation in the abundance of each snRNP and snRNP repertoire (snRNPertoire) in a cell-and tissue-specific manner^9-11^. These observations suggest that differential snRNP abundance may play a role in regulating cell-specific gene expression. However, the mechanisms by which cells maintain distinct snRNP levels and regulate snRNPertoire remain elusive. Additionally, mis-regulation of splicing (due to mutations in splicing factors, snRNP-specific proteins^12, 13^, or non-coding snRNAs^14-18^) can alter splicing outcomes, leading to production of protein isoforms linked to pathogenesis. Therefore, maintaining a sufficient and diverse repertoire of snRNPs is crucial for ensuring their proper function in spliceosome assembly and splicing.

All snRNPs contain Sm cores, which consist of a heptameric Sm protein ring surrounding a short RNA sequence known as the Sm site, typically composed of the nucleotide sequence AU_4-6_G. Assembly of the Sm (or Sm-like) core on each snRNA takes place in the cytoplasm and is chaperoned by the SMN complex^19^. The macromolecular SMN complex comprises oligomeric SMN, Gemins2–8, and unrip proteins; categorized into three functional subunits^20, 21^: 1) SMN/Gemin2 pre-assembles five Sm proteins^22-25^; 2) Gemin5/3/4 recognizes an Sm site and 3’-end of snRNAs, the snRNP code^26-29^; 3) Gemin6/7/8/unrip bind the oligomeric SMN protein, crucial for the function of the SMN complex in snRNP biogenesis^25, 30-33^. When snRNAs lacking Sm sites are overexpressed, they cannot form the Sm core or mature snRNPs, as these snRNAs are subsequently degraded by RNA exonucleases^34, 35^. Importantly, SMN protein deficiency causes spinal muscular atrophy (SMA), a degenerative disease affecting motor neurons in infants^36, 37^. The severity of SMA directly correlates with SMN levels and a corresponding deficit in Sm core assembly^38, 39^. Thus, precise regulation of Sm core assembly by the SMN complex is essential for maintaining adequate snRNP abundance and repertoire within cells.

U1 snRNP (U1) is the most abundant spliceosomal snRNP in human cell (HeLa^40^) and plays a pivotal role in regulating co-transcriptional gene expression through two distinct processes in nucleus: splicing and telescripting. In splicing, U1 recognizes pre-mRNAs through U1 snRNA: 5’ splice site (5’ss) base-pairing during the first step of spliceosome assembly^41^. Simultaneously, in telescripting, U1 suppresses premature cleavage and polyadenylation of nascent transcripts from cryptic polyadenylation signals in introns and 3’-untranslated regions^42, 43^. Furthermore, U1 determines mRNA length and transcription directionality from bi-directional promoters^44-48^. Recent studies shown an enrichment of 5’ss motifs in chromatin-associated long non-coding RNAs are enriched with, suggesting a broader role for U1 in shaping transcriptome^49^. U1 comprises of 11 subunits, including a 5’-trimethyl-G-capped U1 snRNA with four stem-loops (SLs), three U1-specific proteins (U1-70K, U1A, and U1C), and a heptameric Sm core^50^. Structural evidence of U1 has provided functional insights into its biogenesis^51-53^. U1-specific RNA binding proteins U1-70K and U1A bind SL1 and SL2 of U1 snRNA, respectively. The zinc-finger domain of U1C associates with U1-70K through its N-terminus domain, and the heptameric Sm proteins bind to the Sm site (AUUUGUG) in U1 snRNA. Although the SMN complex is essential for snRNP biogenesis, it does not fully explain U1’s over-abundance. U1-70K, previously known as a component of mature U1, hijacks the SMN complex and promotes U1 snRNA-specific Sm core assembly^54^. This protein provides a unique pathway within the SMN complex, promoting U1 formation while suppressing other snRNPs.

Here, we found that the spliceosomal U1-specific U1C protein is a key regulator of the SMN complex for not only U1 but also other snRNPs biogenesis. Together with U1-70K, U1C helps U1 formation, which allows the SMN complex to recruit the other snRNAs:Gemin5 subunit for Sm core assembly. Moreover, U1 snRNA mutations identified in numerous cancers impair Sm core assembly on U1 snRNA *in vitro*. Notably, a point mutation near the U1C binding site hinders the SMN complex’s ability to form canonical snRNPs. These findings highlight the crucial role of U1C in regulating snRNPs biogenesis and suggest that cancer-associated mutations in U1 snRNA may sequester the SMN complex, leading to disruptions in broader RNA metabolism.

## RESULTS

### U1-specific U1C is essential for assembly of the Sm core on all snRNAs

Previous studies show that U1-70K bridges U1 snRNA and the SMN complex, providing an additional and U1-exclusive Sm core assembly pathway^54^. Notably, siRNA-mediated knockdown of U1-70K protein has been shown to concomitantly reduce U1C protein expression but not vice versa^55, 56^, suggesting an interdependent relationship between these proteins. To determine whether U1C plays a role in snRNP biogenesis, we employed the *in vitro* Sm core assembly assay, a high-throughput and quantitative method for assessing Sm core assembled snRNA levels using anti-Sm antibodies^39^. First, HeLa cell extracts were used in which U1-specific proteins (U1-70K and U1C) and SMN complex proteins (SMN and Gemin5) had been knocked down by RNA interference (RNAi). Compared to control RNAi transfection, target-specific siRNAs resulted in a 70–90% reduction in protein level in the cytoplasmic fraction, as confirmed by western blot (WB) (**Figure 1A**). In the assays, *in vitro* transcribed and biotinylated precursor U1 snRNA (pre-U1), other snRNAs (pre-U2, pre-U4, and pre-U5), or a negative control (an Sm site mutant, ΔSm^19^) were incubated in cytoplasmic extracts, respectively. Subsequently, ATP-dependent Sm core-assembled snRNAs were quantified by chemiluminescence.

**Figure 1.**
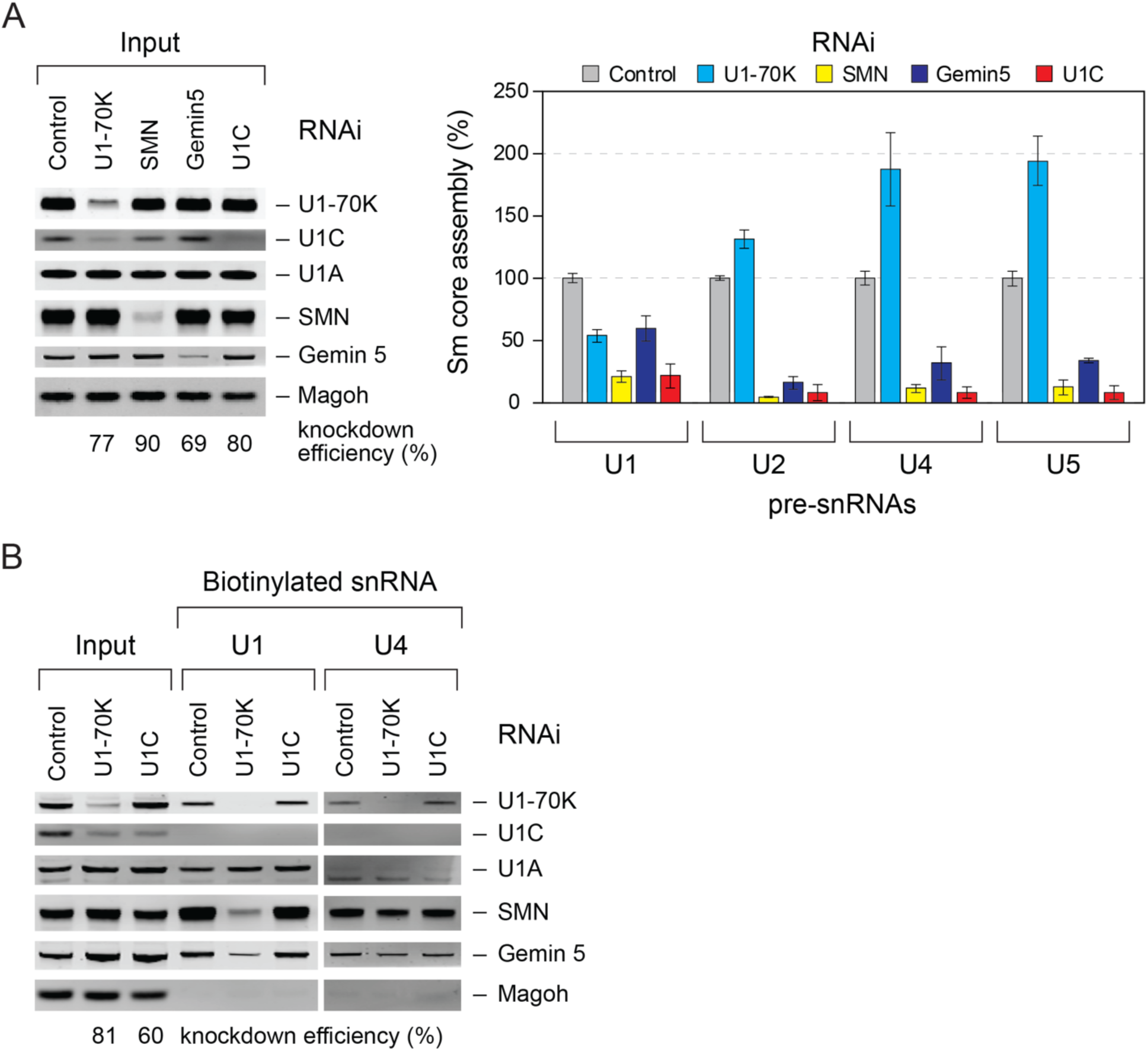
U1C facilitates Sm core assembly on all spliceosomal snRNAs and releases U1 snRNP from the SMN complex *in vitro*. (A) Quantitative western blot analysis of the cytoplasmic extracts from HeLa cells with control, U1-70K, SMN, Gemin5, or U1C short interference RNA (siRNA) knockdown. The knockdown efficiencies relative to Magoh protein as a loading control are indicated as percentages of each protein knockdown compared with control (set at 100%). The Sm core assembly activities on each snRNA in the same cell extracts are compared with control RNAi extracts (100% activity). Error bars represent the standard deviation (SD) from three independent cell cultures. (B) Western blot analysis of SMN complex proteins bound to biotinylated pre-U1 and pre-U4 snRNAs in HeLa cells with control, U1-70K, or U1C siRNA knockdown in the presence of ATP. The input lanes show 20% of each of the cell extracts used. The knockdown efficiencies relative to Magoh protein are indicated. See also Figures S1 and S2.

As expected, knockdown of U1-70K resulted in a 50% reduction in Sm core assembly activity for U1 snRNA, whereas other snRNAs showed a 30–100% increase (**Figures 1A and S1**). Gemin5 knockdown significantly decreased Sm core assembly on other snRNAs by 65–85%, whereas it reduced assembly on U1 by less than 40%. SMN knockdown abolished Sm core assembly on all snRNAs, highlighting its pivotal role in this process. Strikingly, U1C knockdown significantly reduced Sm core assembly not only on U1 but also on other snRNAs, similar to the effect of SMN knockdown. Overexpression of U1-specific proteins in HEK293T cells differentially modulated Sm core assembly on all snRNAs *in vitro* (**Figure S2**). Specifically, U1-70K or U1A facilitated Sm core assembly on U1 snRNA, whereas U1-70K or U1C promoted the assembly on other snRNAs. Thus, these findings indicate a new role for U1C in Sm core assembly across spliceosomal snRNAs, independent of U1-70K and U1A. Our results suggest that U1C has a broader function in regulating the SMN complex and snRNP biogenesis beyond its role as a component of U1.

### U1C is necessary for the release of Sm core-assembled U1 snRNP from the SMN complex

To examine whether U1C influences the interaction between snRNAs and the SMN complex, we performed *in vitro* snRNA pull-down experiments. Binding experiments were performed at 30°C in the presence of ATP to capture intermediate complexes of SMN complex and snRNP-specific proteins on biotinylated snRNAs during Sm core assembly. Following knockdown of HeLa cells with specific siRNAs, cytoplasmic extracts were incubated with snRNAs, and resulting snRNA-protein complexes were isolated with streptavidin beads and analyzed by WB (**Figure 1B**). Notably, U1-70K knockdown abolished U1 snRNA binding to the SMN complex, while U4 snRNA binding was unaffected. In contrast, U1C knockdown maintained the binding of U1 and U4 snRNAs to the SMN complex compared with the control knockdown. Under mild washing conditions (200 mM NaCl), U1C did not tightly bind to U1 snRNA in control, likely due to its lack of an RNA binding domain. Instead, it interacted indirectly with the Sm core through the N-terminus of U1-70K^57, 58^. Strikingly, U1C knockdown did not disrupt the interaction between the SMN complex and pre-U1 or pre-U4 snRNA, although it completely prevented Sm core assembly (**Figure 1B**). Notably, pre-U4 snRNA bound to U1-70K, not U1A, suggesting that the interaction is not RNA-mediated but instead occurs through the oligomeric SMN associated with U1-70K. These results indicate that U1C facilitate the release of Sm core-assembled U1 from the SMN complex, thereby indirectly influencing the assembly of other snRNAs.

Next, we performed immunoprecipitation experiments to determine whether U1C knockdown had any effects on Gemin5’s association with other snRNAs, a key intermediate delivered to the SMN/Gemin2 subunit for Sm core formation^29^. To capture transiently associated snRNAs with Gemin5, HeLa cells transfected with U1-70K or U1C RNAi were crosslinked using 0.2% formaldehyde and immunoprecipitated with anti-Gemim5 antibody. After RNA-Protein complex elution and reversal of crosslinking, we analyzed snRNA abundance using real-time quantitative PCR (RT–qPCR) (**Figure 2A**). U1-70K knockdown increased U1 and U4 snRNA binding with Gemin5 by 4- to 8-fold compared with control knockdown. On the other hand, U2 snRNA binding decreased by half, whereas U5 snRNA remained unaffected. Notably, U1C knockdown increased all tested snRNAs binding to Gemin5 by 4- to 5-fold compared with control knockdown. These results indicate despite U1C and U1-70K being the key components of U1 snRNP, these two proteins showed distinct snRNAs association with Gemin5. The differential bindings of snRNAs to Gemin5 also support the previous observation that at least two distinct snRNA binding sites exist within the SMN complex. One site shows high affinity for U1 and U4 snRNAs, while another has reduced affinity for U2 and U5 snRNAs^26^. Thus, these findings demonstrate that U1C is essential for both the formation and release of mature U1 snRNP from the SMN complex. Furthermore, U1C influences other snRNAs’ access to the SMN complex through Gemin5.

**Figure 2.**
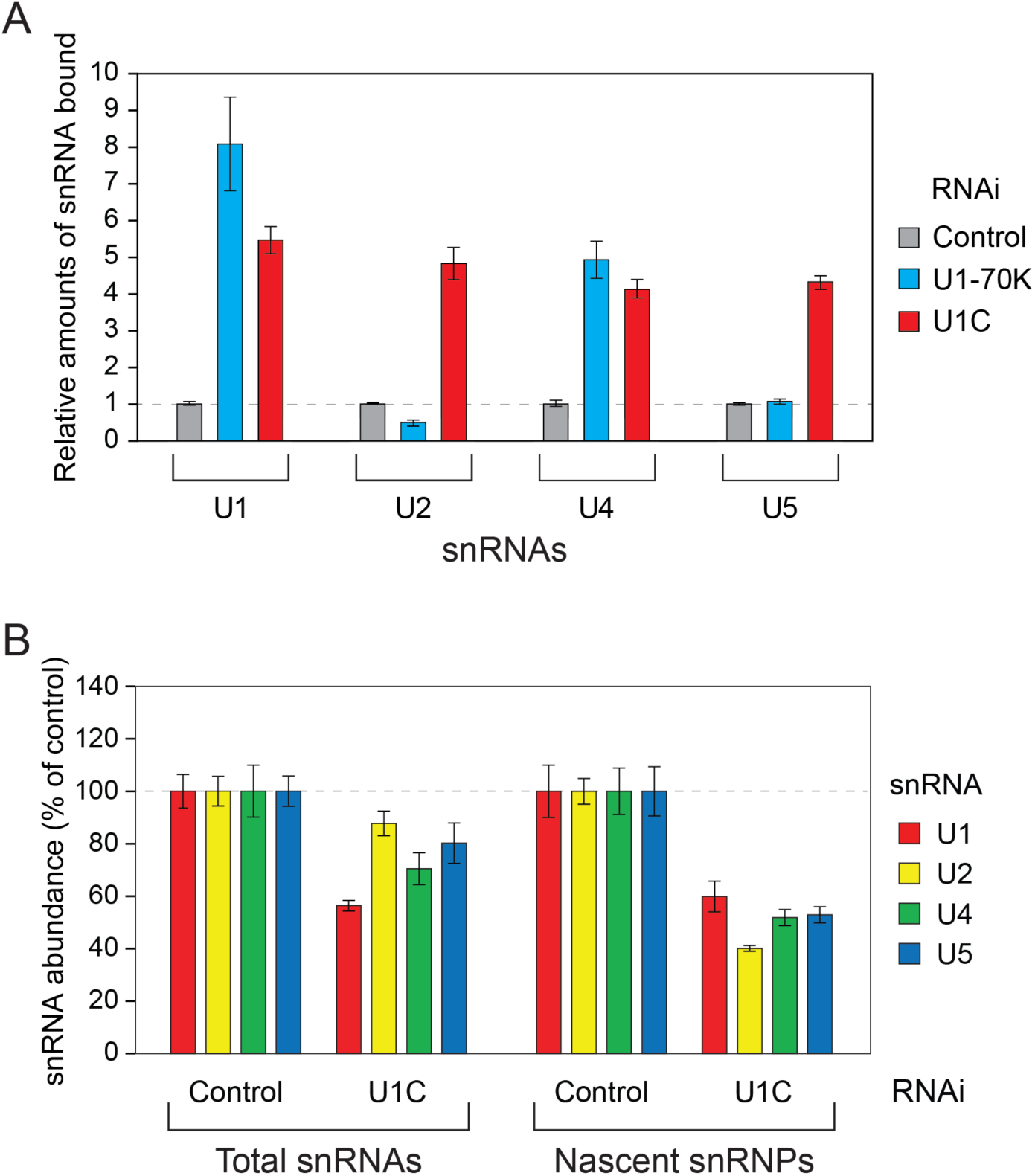
U1C regulates the association of U1 and other snRNAs with Gemin5, as well as the abundance of snRNPs in cells. (A) Quantitative measurement of snRNAs associate with Gemin5 in HeLa cells with control, U1-70K, or U1C siRNA knockdown by quantitative real-time PCR (qRT–PCR). Immunoprecipitations were performed using anti-Gemin5 in cells crosslinked with 0.2% formaldehyde. The relative amounts of each snRNAs compared with control knockdown set are shown. Error bars represent the SD from two independent cell cultures. (B) Quantitative measurements of snRNAs from total RNAs and nascent snRNPs in HeLa cells with control or U1C knockdown. The percent of snRNA abundance compared with those under control RNAi are shown. Error bars represent the SD of three replicates. See also Table S1.

### U1C regulates the snRNP abundance in cells

Previous studies have shown that U1-70K interacts with the SMN complex, facilitating U1 formation while inhibiting the assembly of other snRNPs^54^, thus contributing to U1’s relative abundance in cells. To evaluate U1’s role in regulating snRNP levels, we examined the effect of U1C knockdown on snRNP abundance in cells. U1C knockdown decreased total snRNA levels by 10–40% compared to control knockdown (**Figure 2B**, total snRNAs). Additionally, nascent snRNPs in HeLa cells were quantified by qRT–PCR following anti-Sm immunopurification and metabolically pulse-labeled 4-thiouridine selection. Consistent with *in vitro* Sm core assembly results, U1C knockdown reduced the abundance of all snRNPs by 30–40% compared with control knockdown (**Figure 2B**, nascent snRNPs). These findings indicate that U1C is essential for U1 snRNP formation and also contributes to the formation of other snRNPs in cells.

### U1C interacts with SMN and U1-70K via its two distinct domains

We hypothesized that the reduced Sm core assembly upon U1C knockdown *in vitro* might be due to a potential loss of interaction between U1C with the SMN complex. Several biochemical and structural studies on the U1 snRNP support the role of U1C in stabilizing the mature U1 snRNP through its N-terminal domain (NTD) interactions with U1-70K and Sm proteins^51-53, 59^. However, the function of the C-terminal domain (CTD) of U1C in snRNP biogenesis remains unclear as it is not visible in the human U1 snRNP structure. To test whether U1C directly interacts with the SMN complex, we performed immunoprecipitation experiments in cytoplasmic HeLa cell extracts using anti-U1-70K, anti-U1C, and anti-U1A antibodies, respectively. Subsequent WB analysis revealed that all U1-specific proteins bind to SMN proteins, with U1-70K exhibiting the strongest interaction, followed by U1A and U1C (**Figure 3A**). SMN and Gemin5 interacted with U1-70K and U1A, but not U1C (**Figure S3**).

**Figure 3.**
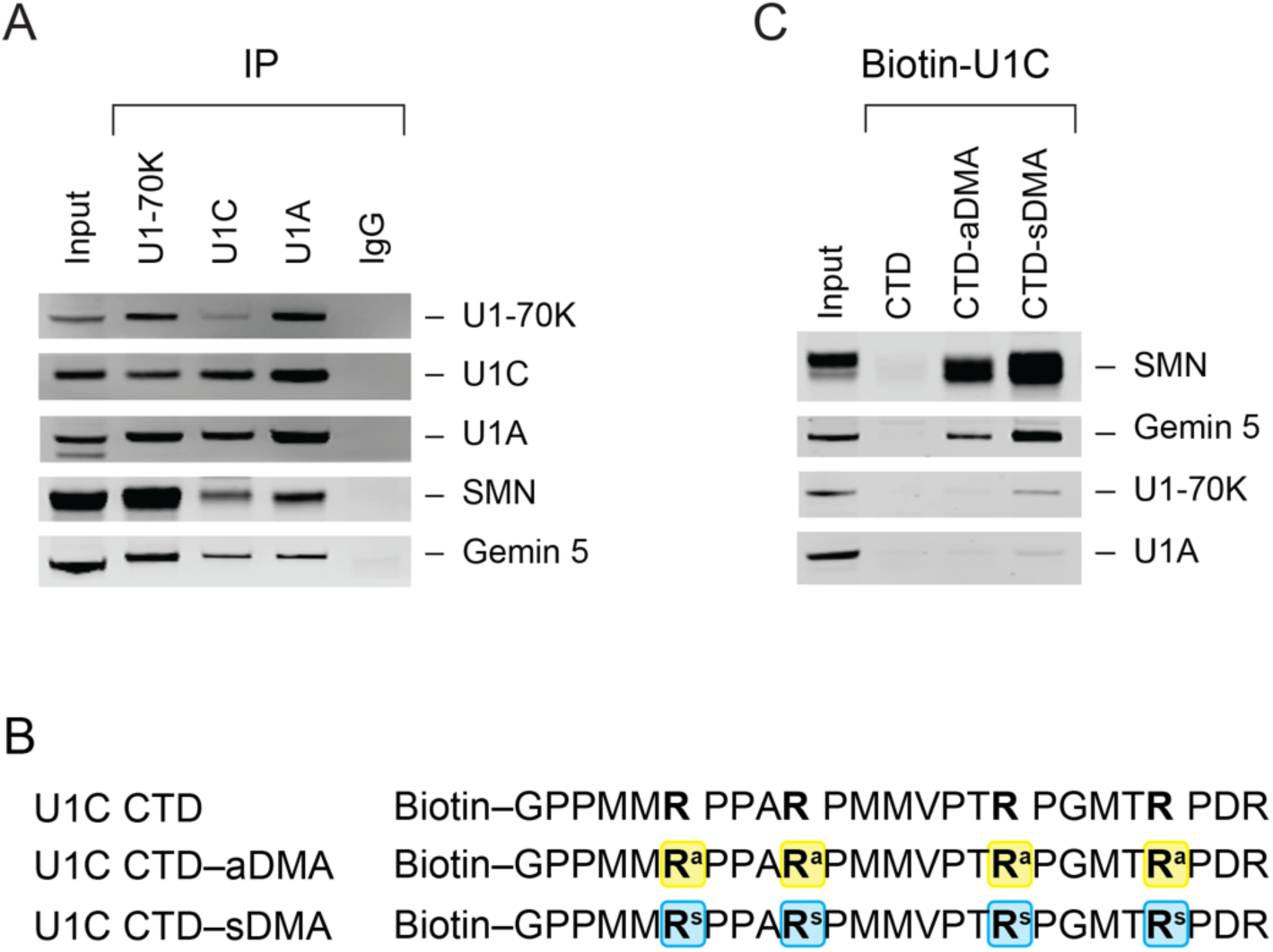
U1C associates with SMN, and this interaction is enhanced by arginine methylation of the U1C C-terminus domain. (A) Western blot (WB) analysis of immunoprecipitated U1 snRNP-specific proteins from cytoplasmic HeLa cell extracts using anti-U1-70K, anti-U1C, anti-U1A antibodies, and mouse immunoglobulin G (IgG). The associated U1-specific proteins, SMN, and Gemin5 are indicated. The input panel shows 2.5% of the proteins used for binding. (B) Primary sequence of the U1C peptide with C-terminus domain (aa 135–159) are shown; U1C CTD (not methylated), U1C CTD–aDMA (asymmetrically demethylated arginine), and U1C CTD–sDMA (symmetrically methylated arginine). Modified arginine residues are highlighted with a box. (C) *In vitro* binding assays using biotinylated U1C CTD peptide with or without DMA in cytoplasmic HeLa extracts were performed, with peptides immobilized on streptavidin beads for binding. The input panel shows 2.6% of the total proteins used. See also Figure S3.

Previous studies have shown that U1C’s CTD is post-translationally modified to di-methylated arginines (DMAs) by CARM1/PRMT4 or PRMT5 methyltransferases^60^. To test whether these modifications enhance the interaction between U1C and SMN protein, we utilized synthetic peptides (25-mer) containing the proline glycine methionine-rich motifs of U1C CTD (aa 135–159) with a 5’-end biotin (**Figure 3B**). These peptides were designed with symmetric (sDMA), asymmetric (aDMA) dimethylated arginines, or no modifications. We performed *in vitro* binding assays using these biotinylated U1C peptides, incubating them in cytoplasmic HeLa cell extracts. The associated proteins were then eluted and analyzed by WB (**Figure 3C**). We observed a robust association between SMN and the sDMA- or aDMA-modified U1C peptide, with a preference for sDMA. Other components of the SMN complex (e.g. Gemin5) were also associated with U1C. However, U1-70K and U1A did not interact with U1C CTD. Notably, the unmodified U1C peptide did not bind the SMN complex. These findings support that DMA modifications of U1C enhance its interaction with the SMN protein, which contains a Tudor domain known to bind proteins with methylated arginine or lysine residues^61-63^. Additionally, these results suggest that U1C’s divalent binding to U1-70K and SMN proteins—through its NTD and CTD, respectively—plays a role in ensuring U1 snRNP formation.

### Mutations in U1 snRNA in various cancers disrupt U1 snRNP formation

Recent studies have identified somatic mutations in U1 snRNA across multiple cancer types, suggesting potential alterations in pre-mRNA splicing^15, 16^. Interestingly, those mutations were distributed throughout the U1 snRNAs transcript, raising questions about the stability and function of U1 snRNP containing these mutations (**Figure 4A**). To investigate the impact of U1 snRNA mutations on Sm core assembly and snRNP formation, we selected several mutations previously reported in 240 cancer patients, spanning 30 different cancer types^15^. We focused on the A3C and A3G mutations in U1 snRNA; the most frequent mutations found in medulloblastoma, chronic lymphocytic leukemia (CLL), and hepatocellular carcinoma (HCC). These mutations are located at the 5’-end of U1 snRNA, a region critical for 5’ splice site recognition^64, 65^. Interestingly, the A3C mutation reduced Sm core assembly by 70% compared to the wild-type (WT), while the A3G mutation had only a minor effect, reducing assembly by 13% (**Figure 4B**). Additionally, other mutations in SLs or 4-way helical junction (4HJ), such as C25T (SL1), C69T (SL2), C46T (4HJ), and C144T (SL4), which are binding sites for U1-specific proteins U1-70K, U1A, as well as U2-specific SF3A1^66^, or are crucial for U1 snRNA structure^67^, reduced Sm core assembly by 40–60%. As expected, the ΔSm mutation, which replaces AUUUGUG to CUCGAG, completely abolished Sm core formation. Moreover, double mutations in A3C and other SL or 4HJ regions, reported in some CLL, B-cell non-Hodgkin lymphoma (B-NHL), and HCC patients, exhibited a more pronounced effect on Sm core assembly defects compared with single mutations (**Figure S4A**). Further analysis of additional 5’-end U1 snRNA mutations (C4G, A7G, C9T, T10A, and G12A) revealed similar, albeit less severe, reductions in Sm core assembly (20–50%) (**Figure S4B**). Given the proximity of U1C to the 5’-end of U1 snRNA in the mature snRNP, we performed *in vitro* Sm core assembly assays using anti-U1C antibodies and observed a correlation between Sm core formation and U1C association (**Figure S4C**). Notably, the C25T mutation in SL1, the binding site for U1-70K, significantly reduced U1C association during Sm core assembly, consistent with previous findings that the U1-70K:Sm core complex creates a binding site for U1C^50, 68^. Our results further suggests that U1-70K binding to U1 snRNA is a prerequisite for U1C association during Sm core assembly on U1 snRNA.

**Figure 4.**
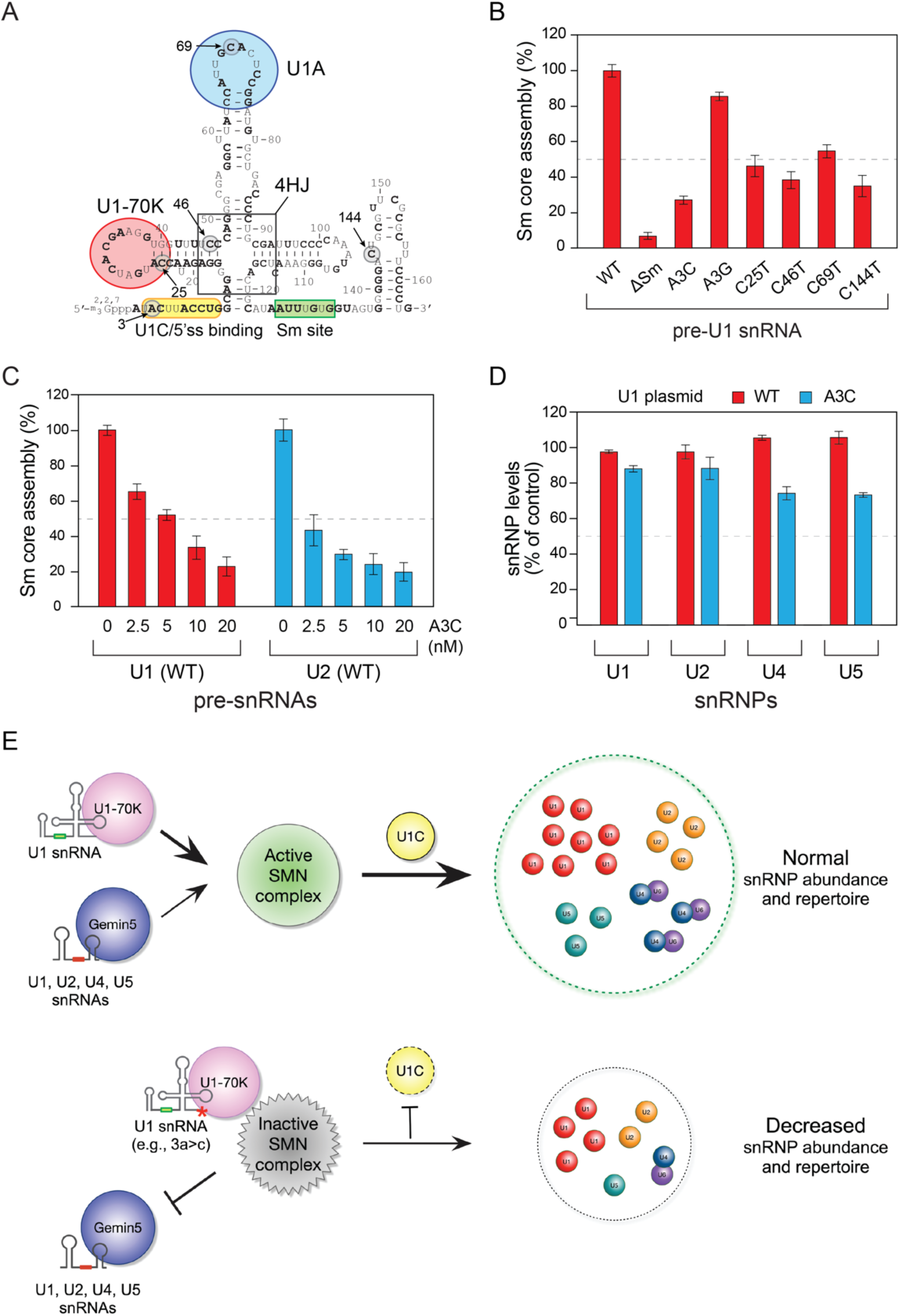
U1 snRNAs mutations identified in cancers disrupt Sm core assembly *in vitro* and alter snRNP abundance in cells. (A) Secondary structure of U1 snRNA, with binding sites for U1-70K, U1A, and U1C/5’ss binding indicated by circles. The Sm site and 4-way helical junction are highlighted in boxes. Residues mutated in pan-cancer patients (Shuai et al., 2019) are shown in bold, and residues tested in this study are marked with arrows. (B) The Sm core assembly activity of U1 snRNA mutants (A3C, C25T, C46T, C69T, C144T) compared to WT, set at 100%. ΔSm serves as a negative control. (C) A3C inhibits Sm core assembly on WT U1 and U2 snRNAs in a dose-dependent manner. Sm core assembly assays were performed with increasing amounts of the non-biotinylated A3C U1 snRNA (2.5–20 nM) and 10 nM biotinylated U1 or U2 snRNA. Error bars represent the SD from three independent cell cultures. (D) *In vivo* snRNP abundance in HEK293T cells transfected with WT, A3C or control plasmid was measured by anti-Sm immunopurification and qRT–PCR. The percentage of snRNA abundance in WT or A3C-transfected cells, relative to control plasmid (set at 100%), is shown. Error bars represent the SD from three replicates. **(E)** Schematic representation of U1C’s role as a gatekeeper for the SMN complex in snRNP biogenesis. Top: U1C associates with a U1 snRNP intermediate, including U1-70K:U1 snRNA, organized within the SMN complex to facilitate U1 snRNP formation. U1C-assisted U1 snRNP formation enables the SMN complex to recruit Gemin5 and other snRNAs (U2, U4, or U5). Bottom: A deficiency in U1C binding to 5’ end U1 snRNA mutants (e.g., 3a>c) sequesters the SMN complex and U1-70K:U1 snRNA, disrupting Sm core assembly on canonical snRNAs. See also Tables S1, S2, and Figure S4.

Since both the SMN complex and U1-70K recognize U1 snRNA, we hypothesized that A3C might share a common binding site on these proteins. To investigate the impact of A3C on Sm core assembly of U1 and other snRNAs, we performed competition Sm core assembly assays. Biotinylated WT U1 snRNA was incubated with cytoplasmic extracts containing various concentrations (2.5–20 nM) of unlabeled A3C. A3C inhibited WT U1 snRNA for Sm core assembly and showed more effective competition with U2 snRNA at equal concentrations (10 nM; **Figure 4C**). To assess the impact of A3C expression on mature snRNP levels in cells, we purified snRNPs using anti-Sm antibodies from HEK293T cells transiently transfected with WT or A3C U1 snRNA plasmids and quantified snRNA abundance using qRT–PCR. Consistent with *in vitro* results, over-expression of WT U1 snRNA plasmid resulted in similar or slightly increased snRNP abundance, whereas that of the A3C plasmid gradually reduced the abundance of all snRNPs (**Figures 4D and S4D**). These findings suggest that A3C binds to overlapping sites on the SMN complex and U1-70K, potentially inhibiting Sm core assembly of both U1 and other snRNAs. Consequently, U1 snRNA mutations identified in cancer patients^15^ disrupt Sm core assembly on U1 snRNA by impairing binding of U1-specific proteins and may also sequester the SMN complex, hindering the formation of canonical snRNPs.

## DISCUSSION

Previous studies have demonstrated that the multi-component SMN complex plays a crucial role in snRNP biogenesis, facilitating Sm core formation on snRNAs^20, 21^. However, the mechanisms regulating its chaperone activity remained an open question. Whereas U1-70K and U1C were known as components of mature U1, our results revealed broader and distinct roles for U1C in snRNP biogenesis. Together with U1-70K, U1C facilitates formation of both U1 and other snRNPs by utilizing the SMN complex in a substrate-assisted manner. Based on the U1 structures^51-53^, our biochemical data suggest a stepwise formation of U1 and other snRNP mediated by the SMN complex. U1-70K promotes the Sm core assembly of U1 by bridging between U1 snRNA and the SMN complex via its U1-specific RNA binding domain^54^. This preferential interaction with U1-70K:U1 snRNA with SMN protein limits the access of other snRNAs to the SMN complex, explaining the relative over-abundance of U1. Pre-assembled U1, involving U1-70K:U1 snRNA and the Sm core, provides an additional interaction site for U1C’s NTD, stabilizing the mature U1. U1C also associates with the SMN protein through its CTD via post-translational arginine modifications, enhancing the interaction with Tudor domain of SMN. In the absence of U1C, the pre-assembled U1-70K:U1 snRNA is unable to dissociate from the SMN complex. This prevents the oligomeric SMN complex from recruiting other snRNAs through Gemin5/3/4, ultimately inhibiting formation of other snRNPs. Thus, U1C plays dual roles in snRNP biogenesis as a gatekeeper of the SMN complex: U1C facilitates completion of mature U1 formation and its release from the SMN complex, while also allowing the SMN complex to recruit other snRNAs via Gemin5/3/4 and promoting snRNP assembly (**Figure 4E**). Other post-translational modifications, such as phosphorylation, are also known to regulate the SMN complex subunits association^69-71^, potentially contributing to the dissociation of mature snRNPs from the SMN complex and increasing the snRNA recruitment for Sm core assembly.

The SMN complex, a well-established RNP-exchange depot, interacts with numerous RNA binding proteins such as like-Sm proteins, sno- or scaRNA binding proteins, and fibrillarin through its oligomeric nature and Tudor domain^54, 72, 73^. Our study demonstrates the broader significance of the SMN complex beyond snRNP biogenesis and SMA, as exemplified by cancer-relevant U1 snRNAs potentially sequestering the SMN complex chaperones (**Figure 4E**). Recent findings linking mutations in snRNAs (U2, U4) or Gemin5 to various developmental and neurodegenerative disorders underscore the SMN complex’s expanded function^14, 17, 18, 74-77^. Combined with engineered U1 snRNAs designed to enhance or suppress 5’ splice sites^78-80^, these results suggest potential applications for U1C in modulating U1’s pre-mRNA binding and snRNP biogenesis. Moreover, structural studies could potentially identify strategies to target U1C for therapeutic interventions in splicing and telescripting.

Frequent mutations in protein-coding genes or dysregulation of pre-mRNAs are major contributors to cancer pathogenesis^81-83^. Recent studies revealed that non-coding RNAs including snRNAs, tRNAs, and lncRNAs are versatile molecules in several cancer models^84-87^. We propose that U1 snRNA mutations can disrupt the homeostasis in snRNP biogenesis, potentially contributing to cancer development. Our data demonstrate that the 5’-end of U1 snRNA is critical for Sm core assembly, and a pan-cancer mutation in this region offer a valuable model to explore the physiological relevance of U1 snRNP in cancer. Although the frequency of these mutations may be low in cancer patients, even a small fraction of U1 snRNA mutants could potentially inhibit SMN complex formation or sequester the SMN complex, leading to impaired biogenesis of not only U1 but also other snRNPs. This in turn could hinder the assembly of active spliceosomes, resulting in a shortage of snRNPs relative to transcriptional output, pre-mRNA processing, or translational machinery^43, 82, 88^. Interestingly, U1 snRNA mutation (3a>g) have been identified in Sonic Hedgehog medulloblastoma; whereas 3a>c mutations are mainly found in CLL and HCC^15, 16^. Although a precise mechanism by which A3C vs A3G mutations differentially affect Sm core assembly remains unclear, atomic-level structural information of U1 with A3G mutation could provide insights into its role in alternative U1 formation and alternative splicing.

## ACKNOWLEDGMENT

We thank Drs. Gideon Dreyfuss, Aaron Hoskins, Manuel Ares, Joshua Mendell, Nicholas Conrad, and Douglas Black for advice and insightful discussions; Drs. Jeffrey Campbell and Zibiao Cao from the North Texas Genomic Center; Mrs Donna Coyle from the University of North Texas Health Care for providing access to instruments. This work is supported by the National Institute of General Medical Sciences of the National Institutes of Health under award number R15GM152936 and by start-up funds the University of Texas Arlington.

## AUTHOR CONTRIBUTIONS

D.M.N., S.M, A.A.K, and B.R.S. conceived and designed the project. D.M.N., S.M., A.A.K., J.K., T.M., P.A., A.A., R.R., M.P., and B.R.S. performed the experiments. K.J., E.S., J.Y. contributed to data analysis and discussion of the results. D.M.N., S.M, A.A.K, B.R.S. wrote the manuscript with input from all authors. B.R.S. was responsible for the project planning and performance.

## Key resources table

| REAGENT or RESOURCE | SOURCE | IDENTIFIER |
| --- | --- | --- |
| **Antibodies** |  |  |
| Smith Antigen (Y12) | Sigma-Aldrich | MABF2793 |
| U1C (4H12) | Sigma-Aldrich | SAB4200188;<br>RRID:AB_10640155 |
| U1-70K (H111) | Synaptic Systems | 203011;<br>RRID:AB_887903 |
| U1-70K (E-4) | Santa Cruz<br>Biotechnology | sc-390988<br>RRID:AB_3083771 |
| U1A | Proteintech | 10212-1-AP;<br>RRID:AB_2239723 |
| SMN (8/SMN) | BD Bioscience | 610646;<br>RRID:AB_397973 |
| Gemin3 (2/DP103/Gemin3) | BD Bioscience | 612152;<br>RRID:AB_399523 |
| Gemin5 | Proteintech | 24897-1-AP;<br>RRID:AB_2879784 |
| Magoh (21B12) | Santa Cruz<br>Biotechnology | sc-56724;<br>RRID:AB_629914 |
| Mouse Immunoglobulin G | Sigma-Aldrich | I5381 |
| IRDye 800CW Goat anti-Mouse IgG | Licor Biotech | 92632210 |
| IRDye 800CW Goat anti-Rabbit IgG | Licor Biotech | 92632211 |
| IRDye 800CW Goat anti-Rat IgG | Licor Biotech | 92532219 |
| Fluorescent TrueBlot Anti-Mouse Ig DyLight™ 800 | Rockland<br>Immunochemicals | 18-4517-32 |
| **Bacterial and virus strains** |  |  |
| BL21(DE3)pLysS Competent Cells | Invitrogen | C606010 |
| XL-10 Gold Ultracompetent cells | Agilent | 200314 |
| DH5α Competent Cells | Invitrogen | 18258012 |
| **Biological samples** |  |  |
| **Chemicals, peptides, and recombinant proteins** |  |  |
| UltraPure DNase/RNase-Free Distilled Water | Invitrogen | 10977015 |
| Dulbecco's Modified Eagle Medium (DMEM) | Gibco | 11995065 |
| Opti-Minimum Essential Medium (MEM) | Gibco | 31985062 |
| Fetal Bovine Serum | Sigma-Aldrich | TMS013B |
| L-glutamine | Gibco | 25030081 |
| Penicillin-Streptomycin | Gibco | 10378016 |
| Tris-HCl, 1M solution, pH 7.5 | Invitrogen | 15567027 |
| MgCl<sub>2</sub>, 1M solution | Ambion | AM9530G |
| Ethylenediaminetetraacetic Acid (EDTA), 0.5M solution, pH 8.0 | Ambion | AM9260G |
| Sodium dodecyl sulfate (SDS) | Fisher Scientific | BP8200100 |
| Sodium chloride, 5M solution | Ambion | AM9760G |
| Phenol:Chloroform:isopropylalcohol (25:24:1, v/v) | Fisher Scientific | BP1752 |
| Ethanol, Absolute | Fisher Scientific | BP28184 |
| TRIzol | Invitrogen | 15596026 |
| Formaldehyde (37%, w/v) | Fisher Scientific | BP531 |
| Empigen BB (~30% w/v) | Sigma-Aldrich | 30326 |
| NP-40 alternative | Calbiochem | 492016 |
| Triton X-100 | Fisher Scientific | BP151 |
| Heparin Sodium | Fisher Scientific | BP2425 |
| Digitonin | Calbiochem | 300410 |
| tRNA, from *Escherichia coli* MRE600 | Roche | 10109541001 |
| RNasin Plus Ribonuclease Inhibitor | Promega | N2615 |
| 4-thiouridine | Sigma-Aldrich | T4509 |
| Biotin-16-UTP | Roche | 11388908910 |
| cOmplete™ Protease Inhibitor Cocktail | Roche | 11697498001 |
| Avidin, NeutrAvidin™, Horseradish Peroxidase conjugate | Invitrogen | A2664 |
| SuperSignal™ ELISA Femto Substrate | Thermo Scientific | 37075 |
| ATP solution (100 mM) | Thermo Scientific | R0441 |
| EZ-Link HPDP-Biotin | Thermo Scientific | 21341 |
| Dynabeads MyOne Streptavidin C1 | Invitrogen | 65002 |
| Dynabeads M-280 Streptavidin | Invitrogen | 11206D |
| Dynabeads Protein G | Invitrogen | 10004D |
| NuPAGE LDS Sample Buffer (4X) | Invitrogen | NP0008 |
| Biotinylated-U1C CTD peptide (aa 135–159) without modification:<br>Biotin- GPPMMRPPARPMMPVTRPGMTRPDR -OH | This study, Sigma-Aldrich | N/A |
| Biotinylated-U1C CTD peptide (aa 135–159) with asymmetric arginine di-methylations:<br>Biotin- GPPMM-R(Me)2AS-PPA-R(Me)2AS-PMMVPT-R(Me)2AS-PGMT-R(Me)2AS-PDR -OH | This study, Sigma-Aldrich | N/A |
| Biotinylated-U1C CTD peptide (aa 135–159) with symmetric arginine di-methylations:<br>Biotin- GPPMM-R(Me)2S-PPA-R(Me)2S-PMMVPT-R(Me)2S-PGMT-R(Me)2S-PDR -OH | This study, Sigma-Aldrich | N/A |
| Critical commercial assays |  |  |
| Bradford Protein Assay Kit | Pierce | PI23200 |
| Phusion™ Plus PCR Master Mix | Thermo Scientific | F631XL |
| QuikChange Lightning Site-directed Mutagenesis Kit | Agilent | 210518 |
| MEGAscript™ T7 Transcription Kit | Invitrogen | AM1334 |
| Lipofectamine RNAiMax | Invitrogen | 13778075 |
| SuperScript™ IV First-Strand Synthesis System | Invitrogen | 18091050 |
| PowerTrack™ SYBR Green Master Mix | Applied Biosystems | A46110 |
| Deposited data |  |  |
| Experimental models: Cell lines |  |  |
| Human: HeLa cells | ATCC | CCL-2.2 |
| Human: HEK293T | ATCC | CRL-3216 |
| Experimental models: Organisms/strains |  |  |
| Oligonucleotides |  |  |
| ON-TARGETplus Human SNRPC siRNA | Dharmacon, Horizon discovery | L-019574-02-0050 |
| ON-TARGETplus Human SNRNP70 siRNA | Dharmacon, Horizon discovery | LQ-018811-00-0005 |
| ON-TARGETplus Human SMN1 siRNA | Dharmacon, Horizon discovery | LQ-011108-00-0005 |
| ON-TARGETplus Human GEMIN5 siRNA | Dharmacon, Horizon discovery | LQ-013954-01-0005 |
| Primers for snRNA plasmid construction, see Table S1 | This study | N/A |
| Primers for Site-directed mutagenesis, see Table S2 | This study | N/A |
| Primers for RT-PCR, see Table S3 | Zhang et al. <sup>11</sup> | N/A |
| Recombinant DNA |  |  |
| pUC19 pre-U1 snRNA | Yong et al., <sup>29</sup> | N/A |
| pUC19 pre-U1 snRNA (ΔSm) | Yong et al., <sup>29</sup> | N/A |
| pUC19 pre-U1 snRNA<sup>A3C</sup> | This study | N/A |
| pUC19 pre-U1 snRNA<sup>C25T</sup> | This study | N/A |
| pUC19 pre-U1 snRNA<sup>C46T</sup> | This study | N/A |
| pUC19 pre-U1 snRNA<sup>C69T</sup> | This study | N/A |
| pUC19 pre-U1 snRNA<sup>C144T</sup> | This study | N/A |
| pUC19 pre-U1 snRNA<sup>A3C/C25T</sup> | This study | N/A |
| pUC19 pre-U1 snRNA<sup>A3C/C46T</sup> | This study | N/A |
| pUC19 pre-U1 snRNA<sup>A3C/C69T</sup> | This study | N/A |
| pUC19 pre-U1 snRNA<sup>A3C/C144T</sup> | This study | N/A |
| pUC19 pre-U2 snRNA | This study | N/A |
| pUC19 pre-U2 snRNA (ΔSm) | This study | N/A |
| pUC19 pre-U4 snRNA | This study | N/A |
| pUC19 pre-U4 snRNA (ΔSm) | This study | N/A |
| pUC19 pre-U5 snRNA | This study | N/A |
| pUC19 pre-U5 snRNA (ΔSm) | This study | N/A |
| pBlueScript U1 snRNA | Oh et al., <sup>87</sup> | N/A |
| pBlueScript U1 snRNA<sup>A3C</sup> | This study | N/A |
| Software and algorithms |  |  |
| Protein arginine methylation prediction tool (PRmePRed) | Kumar et al., <sup>89</sup> |  |
| Other |  |  |

## MATERIALS AND METHODS

### Cell culture, transfection, and formaldehyde crosslinking

HeLa and HEK293T cells were cultured in DMEM supplemented with 10% (v/v) fetal bovine serum (FBS), 2 mM L-glutamine, 10 units/mL penicillin, and 10 μg/mL streptomycin at 37°C with 5% CO_2_. HeLa cells were transfected with control siRNA or siRNA targeting U1C, U1-70K, SMN, and Gemin5 (Dharmacon, Horizon Discovery) using RNAiMax (Invitrogen). After 42–46 h of incubation, cells were harvested for snRNA measurements, *in vitro* Sm core assembly, or RNA pull-down assays.

U1 snRNAs in pBlueScript plasmid were transfected in HEK293T cells using the NEON transfection system (Invitrogen) via electroporation of 5×10^6^ cells with two pulses of 20 ms each at 1,100 V. Following electroporation, cells were incubated in DMEM in a 15 cm plate. After 42–48 h of incubation, cells were harvested and washed twice with ice-cold PBS. For nascent snRNP measurements, HeLa cells were transfected with siRNA and then metabolically labeled with 200 µM of 4-thiouridine during the final 2 h of transfection. To capture Gemin5-associated snRNAs in cells, HeLa cells were fixed with 0.2% (w/v) formaldehyde for 10 min at room temperature with gentle rotation. The reactions were quenched with glycine (pH 7.0) to a final concentration of 150 mM for 10 mins, followed by two washes with ice-cold PBS.

### Construction of precursor snRNAs and *in vitro* transcription

Precursor U2, U4, and U5 snRNAs were amplified from HeLa cell genomic DNA using Phusion Plus PCR Master Mix (ThermoScientific) and subsequently inserted into the pUC19 plasmid between EcoRI and HindIII. Mutations in precursor U1 snRNA were introduced via QuikChange Lightning Site-Directed Mutagenesis Kit (Agilent). All snRNA sequences were verified by Sanger Sequencing (Eurofinsgenetics). Linearized pre-snRNA plasmids with BsaI digestion were used for *in vitro* transcription using MEGAscript T7 Transcription Kit (Invitrogen) in the presence of 1.75 mM biotin-16-UTP (Roche) and 1.875 mM UTP. Transcribed RNAs were purified by electrophoresis on a 6% polyacrylamide gel containing 5 M urea; followed by phenol:chloroform extraction, ethanol precipitation, and resuspension in RNase-free DEPC water. The concentrations of pre-snRNAs were determined by UV absorbance at 260 nm.

### U1C peptides

The 5’-end biotinylated U1C C-terminus domain peptides (aa 135-159) with no methylation, aDMA (asymmetrically di-methylated arginine), and sDMA (symmetrically methylated arginine) modification were identified using the Protein arginine methylation prediction tool (PRmePRed^89^, http://bioinfo.icgeb.res.in/PRmePRed/). Custom-designed peptides were obtained from Sigma–Aldrich. Each peptide was dissolved in UltraPure water (Invitrogen) to prepare a stock solution at 2 mM, and further diluted to 20 μM in a buffer containing 10 mM Tris-HCl (pH 7.5), 50 mM NaCl for *in vitro* binding studies.

### Preparation of HeLa cytoplasmic extracts

With a slightly modified version of the Pellizzoni et al. (2002)^19^ procedure, HeLa S3 cells were resuspended in reconstitution buffer [20 mM HEPES/KOH pH 7.9, 50 mM KCl, 5 mM MgCl_2_, 0.2 mM EDTA] containing 100 µg/mL digitonin and protease inhibitor, then passed through a 25G needle five times on ice. The cell lysate was clarified by centrifugation at 1,500×g for 1 min, and the supernatant was mixed with NP-40 to a final concentration of 0.01%. After a second centrifugation at 9,400×g for 15 min at 4°C, the supernatant was collected, glycerol was added to a final concentration of 5%, and the mixture was stored at –80°C. The concentration of various extracts was determined by the Bradford protein assay (Pierce).

### Antibodies, immunoprecipitation, and quantitative immunoblotting

The following antibodies were used for this study: anti-SMN (BD Bioscience or 2B1), anti-Gemin5 (Proteintech), anti-Gemin3 (BD Bioscience), anti-Sm (Sigma–Aldrich), anti-U1C (Sigma–Aldrich), anti-U1-70K (Synaptic Systems or Santa Cruz Biotechnology), anti-U1A (Proteintech), and anti-Magoh (Santa Cruz Biotechnology).

Immunoprecipitation was performed using anti-U1C, anti-U1-70K, anti-SMN, and anti-Gemin5 antibodies immobilized on Dynabeads Protein G (Invitrogen). The beads were incubated with cytoplasmic extracts (2 mg/mL) for 1 h at 4°C in a 96-well plate. After incubation, the beads were washed five times with RSB-100 buffer [10 mM Tris-HCl (pH 7.5), 100 mM NaCl, 2.5 mM MgCl_2_] containing 0.1% NP-40 using a KingFisher Flex magnetic beads processor (ThermoFisher). The beads were then equilibrated with RSB-150 buffer containing 0.02% NP-40, and isolated RNA-protein complexes were eluted with sample buffer. Cytoplasmic cell extracts (20 µg) from HeLa cells or immunoprecipitated proteins were separated by electrophoresis on 12% SDS–PAGE gels and transferred to nitrocellulose membranes. Quantitative western blotting was performed following the manufacturer’s protocol. The membrane was scanned, and the intensity of the protein bands was analyzed on an Odyssey infrared imaging system.

### *In vitro* RNA pull-down

Cytoplasmic extracts (2 mg/mL) from HeLa cells with control or target protein knockdown were incubated with 10 nM of biotinylated snRNA in reconstitution buffer containing 0.25 mg/mL *E. coli* tRNA and 1.0 U/μL RNase inhibitor in 96-well plates for 1 h at 4°C, with gentle mixing at 650 rpm. M280 Streptavidin Dynabeads (Invitrogen) were added in 100 µL of RSB-150 buffer containing 0.02% Triton X-100, a protease inhibitor tablet (Roche), and 0.2 U/µL RNase inhibitor to the reaction mixture and incubated for an additional hour. The beads were washed five times in RSB-200 buffer containing 0.02% Triton X-100 using a magnetic beads processor. Bound proteins were eluted by boiling in 10 µL of sample buffer, and then analyzed by SDS–PAGE and western blotting.

### Isolation of Gemin5:snRNA complexes and nascent snRNPs in cells

Gemin5-associated snRNAs were isolated by immunopurification using anti-Gemin5 antibodies from HeLa cell extracts (2mg/mL) lysed in RSB-300 buffer, containing 1% (w/v) Empigen BB and 0.5% (w/v) NP-40 for immunoprecipitation. Collected beads were then incubated with 1 mg/mL of final concentration protease K in a buffer [50 mM Tris-HCl (pH 7.5), 150 mM NaCl, 1% SDS, 5 mM EDTA (pH 8.0)] at 30°C for 30 min. The snRNAs were recovered by phenol:chloroform extraction, ethanol precipitation, and resuspension in 20 µL of RNase-free DEPC water. The concentrations of snRNAs were determined by UV absorbance at 260 nm. Total snRNPs were isolated by immunopurification using anti-Sm antibodies from HeLa cell extracts lysed in RSB-500 buffer containing 0.1% NP-40, protease-inhibitor, and 0.2 U/μL RNase inhibitor. After 1 h incubation at 4°C, the beads were washed five times with the lysis buffer using the magnetic beads processor. The same procedures for Gemin5:snRNA complexes purification was applied to isolate snRNAs from the beads. For nascent snRNP purification, snRNAs from total snRNPs were biotinylated with 0.2 mg/mL of EZ-Link HPDP-Biotin and captured using Dynabeads MyOne Streptavidin beads, as previously described^54^.

### Reverse transcription and RT–qPCR measurement of snRNAs

Total RNA was extracted from cells using Trizol (Invitrogen). The specific primers for snRNA, 5S, and 5.8S rRNA were used to generate complementary DNAs (cDNAs) using SuperScript IV Reverse Transcriptase Kit (ThermoFisher). Either 300 ng of total RNA or 1.5 µL of immunopurified RNAs were used as a template in a 20 µL reaction. For each snRNA, 1% (v/v) of the cDNA was used for real-time quantitative PCR (RT-qPCR). PCR reactions (10 µL) were carried out using SYBR Green Master mix (ThermoFisher) and a CFX Duet Real-Time PCR system (Bio-Rad). Each snRNA was measured in triplicate. A plasmid encoding snRNA sequence was used as a standard for absolute quantification of immunoprecipitated snRNAs. Details of all primers and probes are in Table S1.

### *In vitro* Sm core assembly and U1C binding assays

The quantitative, high-throughput Sm core assembly assay was performed in accordance with a previously established protocol^39, 90^. Briefly, cytoplasmic extracts (2 mg/mL) from HeLa cells were incubated with biotinylated snRNAs (10 nM) in reconstitution buffer containing 2.5 mM ATP, 0.25 mg/mL *E. coli* tRNA, and 1.0 U/μL RNase inhibitor for 1 h at 30°C. For competition experiments, biotinylated WT U1 or U2 snRNAs (10 nM) were incubated with increasing concentrations of non-biotinylated pre-U1 mutants (2.5, 5.0, 10.0, and 20.0 nM) in HeLa cytoplasmic extracts for 1 h at 30°C. Sm core assembled snRNPs were captured using anti-Sm antibodies-immobilized protein G magnetic beads in RSB-500 buffer containing 0.1% NP-40, protease inhibitor, 2 mg/mL heparin sulfate, and 0.2 U/μL RNase inhibitor for an additional hour at 30°C. The beads were washed four times with RSB-500 buffer containing 0.1% NP-40, followed by a final wash with RSB-150 buffer containing 0.02% NP-40, using a magnetic particle processor. The snRNPs on the beads were incubated in 0.04 μg/mL of horseradish peroxidase–conjugated Avidin in RSB-150 buffer containing 0.02% NP-40 and incubated for 1 h at 30°C. The beads were then washed as before and resuspended in 150 μL of SuperSignal ELISA Femto substrate (Pierce). Chemiluminescence signals were detected at 495 nm with a BioTek Synergy HTX microplate reader.

For U1C binding assays with U1 snRNAs, a modified Sm core assembly assay was used, with U1C antibody-immobilized beads substituting for anti-Sm antibodies in the procedure. Additionally, in vitro Sm core assembled U1 snRNPs were purified in RSB-50 containing, 0.02% TritonX-100.

### *In vitro* U1C peptide binding assays

Biotinylated U1C peptides (0.5 nmol) were first immobilized on Dynabeads MyOne Streptavidin C1 (Invitrogen) in 100 µL of RSB-200 containing 0.02% Triton X-100 for 30 min. The beads were then incubated with HeLa cytoplasmic extracts (2 mg/mL) for 1 h at 4°C. After incubation, the beads were washed five times with the same buffer using a magnetic particle processor. Bound proteins were eluted by boiling in 25 µL of sample buffer, and then analyzed by SDS–PAGE and western blotting.

## Supplemental Information

**Table S1.**
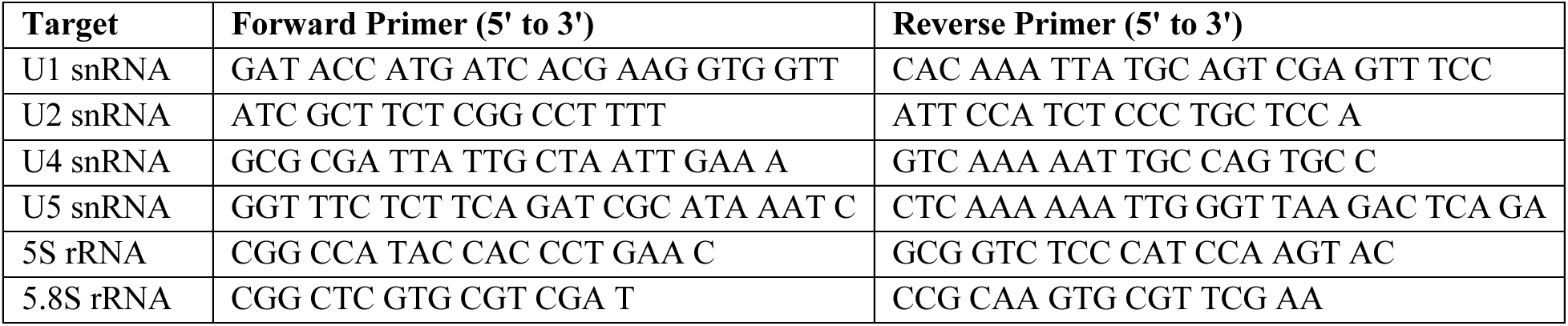
Primers used for RT–qPCR.

**Table S2.**
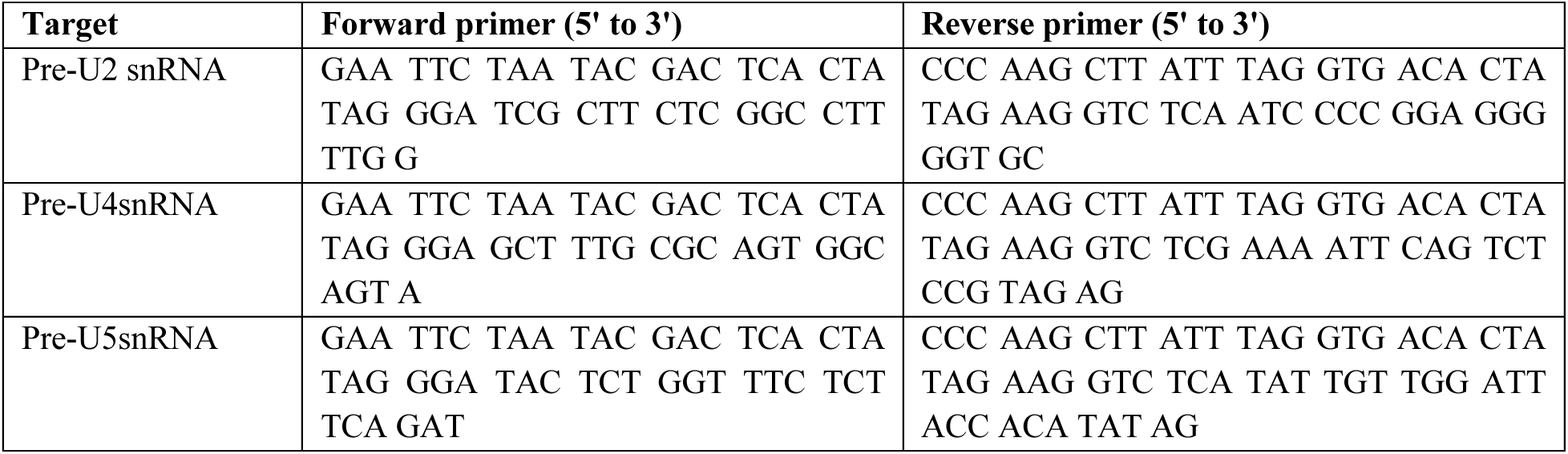
Primers used for snRNA plasmid construction in this study.

**Table S3.**
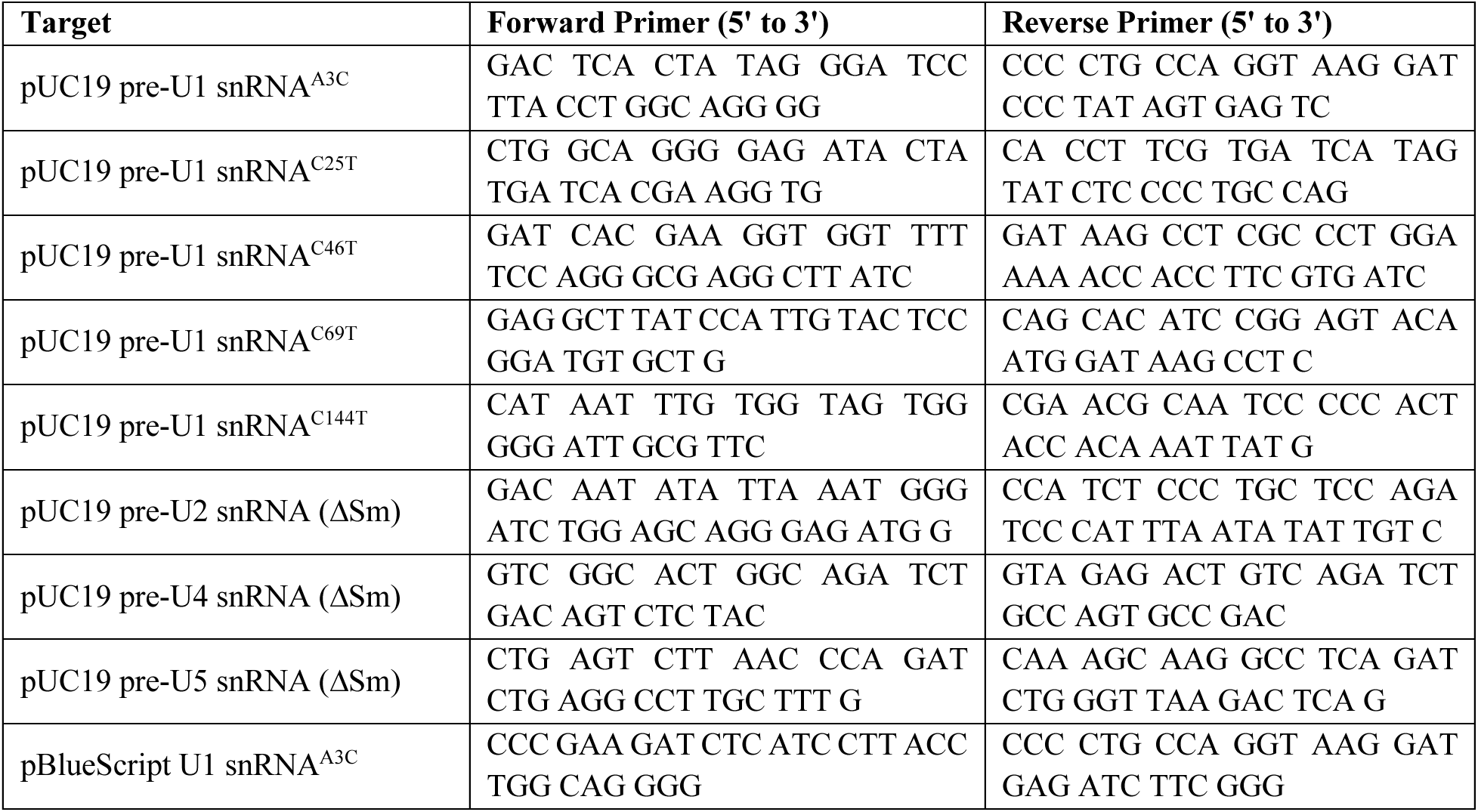
Primers used for site-directed mutagenesis of snRNA in this study.

**Figure S1.**
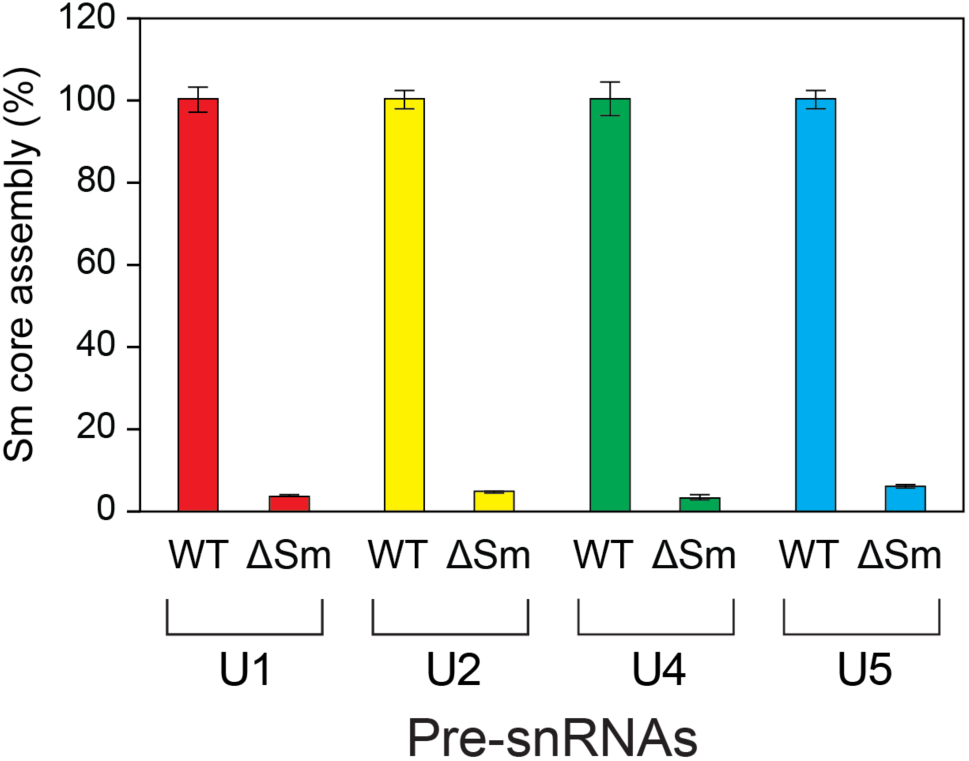
Sm site mutant (ΔSm) pre-snRNAs show defective Sm core assembly compared to wild-type snRNAs (related to. Figure 1). Sm core assembly activity on each pre-snRNA with an Sm site mutant (ΔSm) was compared to wild-type (WT) snRNA, which was set as 100% activity. Error bars represent the SD from three replicates.

**Figure S2.**
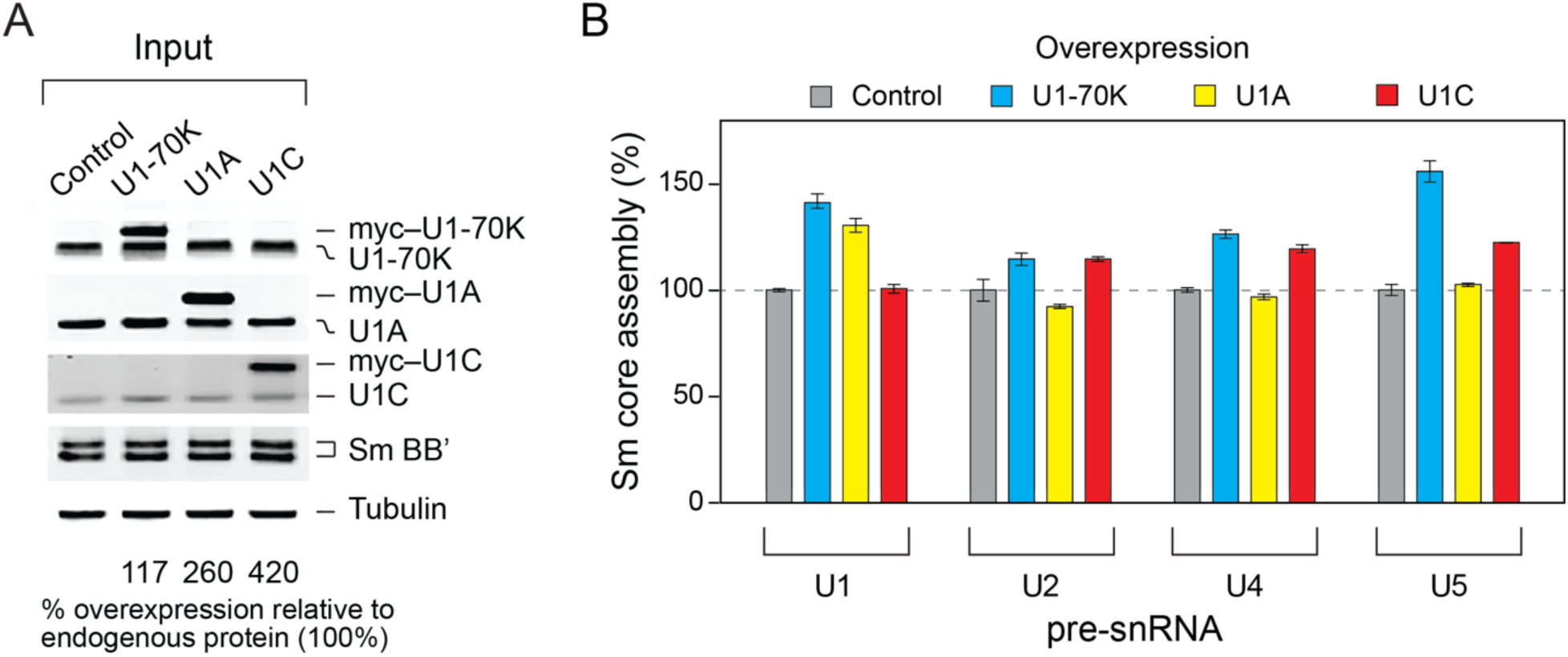
Overexpression of U1-specific proteins lead to differential Sm core assembly across all snRNAs (related to Figure 1). (A) Western blot analysis of overexpressed U1-specific protein (myc–U1-70K, myc–U1A, and myc–U1C) compared to endogenous proteins (set at 100%). Input lanes show 10 µg of each of cytoplasmic HEK293T cell extracts. Sm B/B’ represents Sm proteins and tubulin is used as the loading control. (B) Sm core assembly activities on each snRNA is compared to control extracts (100% activity). Error bars represent SD from three replicates.

**Figure S3.**
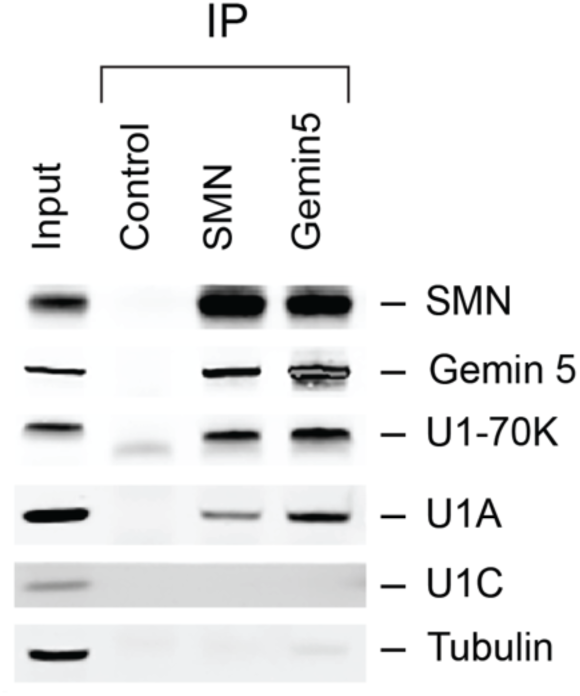
Interaction of SMN and Gemin5 with U1-specific proteins (related to Figure 3). SMN and Gemin5 were immunoprecipitated using target-specific antibodies, followed by SDS-PAGE and western blot analysis using antibodies specific to SMN complex or U1-specific proteins. The input lane shows 2% of the cytoplasmic HeLa extracts used for binding. The control lane shows the background signal from immunoprecipitation with mouse Immunoglobulin G (IgG).

**Figure S4.**
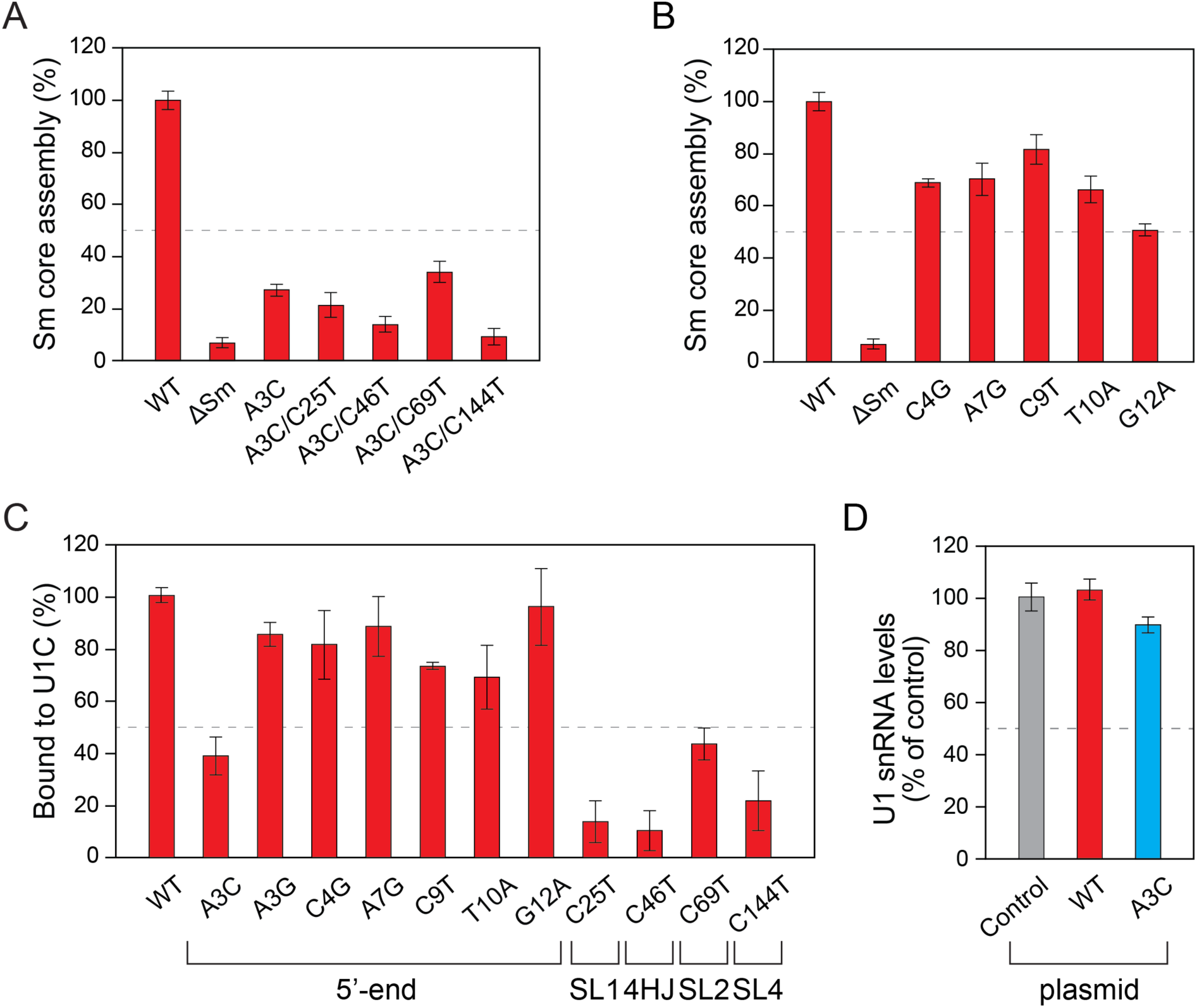
Mutations in U1 snRNA from cancer patients exhibit various effects on in vitro Sm core assembly, U1C binding, and total U1 snRNA expression in cells (related to Figure 4). (A) The Sm core assembly activity of the U1 snRNA mutants (A3C alone and double mutants including A3C/C25T, A3C/C46T, A3C/C69T, and A3C/C144T) in HeLa cells is shown compared with WT, set at 100%. The ΔSm U1 snRNA is a negative control. (B) Same assays as in (A) but with the 5’-end U1 snRNA mutations (C4G, A7G, C9T, T10A, and G12A). (C) In vitro U1C binding experiments with U1 snRNA mutations at the 5’-end, stem loops (SL1, SL2, and SL4), and a 4-way helical junction (4HJ). Error bars represent the SD based on three independent cell cultures. (D) HEK293T cells (5 million) were transfected with 1.5 μg of WT or A3C U1 snRNA in pBlueScript plasmid with native promoter via electroporation and incubated for 48 hr. Isolated total RNAs were treated with DNase then total U1 snRNAs abundance was determined by RT-qPCR. RNA levels were normalized to 5S and 5.8 rRNA. Error bars represent the SD from triplicate samples.

## References

1. Shine, M., Gordon, J., Schärfen, L., Zigackova, D., Herzel, L., and Neugebauer, K.M. (2024). Co-transcriptional gene regulation in eukaryotes and prokaryotes. Nat Rev Mol Cell Biol 25, 534–554.

2. Gehring, N.H., Wahle, E., and Fischer, U. (2017). Deciphering the mRNP Code: RNA-Bound Determinants of Post-Transcriptional Gene Regulation. Trends Biochem Sci 42, 369–382.

3. Patel, A.A., and Steitz, J.A. (2003). Splicing double: insights from the second spliceosome. Nat Rev Mol Cell Biol 4, 960–970.

4. Tian, B., Pan, Z., and Lee, J.Y. (2007). Widespread mRNA polyadenylation events in introns indicate dynamic interplay between polyadenylation and splicing. Genome Res 17, 156–165.

5. Lee, Y., and Rio, D.C. (2015). Mechanisms and Regulation of Alternative Pre-mRNA Splicing. Annu Rev Biochem 84, 291–323.

6. Will, C.L., and Lührmann, R. (2001). Spliceosomal UsnRNP biogenesis, structure and function. Curr Opin Cell Biol 13, 290–301.

7. Shi, Y. (2017). Mechanistic insights into precursor messenger RNA splicing by the spliceosome. Nat Rev Mol Cell Biol 18, 655–670.

8. Fica, S.M., and Nagai, K. (2017). Cryo-electron microscopy snapshots of the spliceosome: structural insights into a dynamic ribonucleoprotein machine. Nat Struct Mol Biol 24, 791–799.

9. Dvinge, H., Guenthoer, J., Porter, P.L., and Bradley, R.K. (2019). RNA components of the spliceosome regulate tissue- and cancer-specific alternative splicing. Genome Res 29, 1591–1604.

10. Mabin, J.W., Lewis, P.W., Brow, D.A., and Dvinge, H. (2021). Human spliceosomal snRNA sequence variants generate variant spliceosomes. RNA 27, 1186–1203.

11. Zhang, Z., Lotti, F., Dittmar, K., Younis, I., Wan, L., Kasim, M., and Dreyfuss, G. (2008). SMN deficiency causes tissue-specific perturbations in the repertoire of snRNAs and widespread defects in splicing. Cell 133, 585–600.

12. Yoshida, K., Sanada, M., Shiraishi, Y., Nowak, D., Nagata, Y., Yamamoto, R., Sato, Y., Sato-Otsubo, A., Kon, A., Nagasaki, M., et al. (2011). Frequent pathway mutations of splicing machinery in myelodysplasia. Nature 478, 64–69.

13. Bradley, R.K., and Anczuków, O. (2023). RNA splicing dysregulation and the hallmarks of cancer. Nat Rev Cancer 23, 135–155.

14. Jia, Y., Mu, J.C., and Ackerman, S.L. (2012). Mutation of a U2 snRNA gene causes global disruption of alternative splicing and neurodegeneration. Cell 148, 296–308.

15. Shuai, S., Suzuki, H., Diaz-Navarro, A., Nadeu, F., Kumar, S.A., Gutierrez-Fernandez, A., Delgado, J., Pinyol, M., López-Otín, C., Puente, X.S., et al. (2019). The U1 spliceosomal RNA is recurrently mutated in multiple cancers. Nature 574, 712–716.

16. Suzuki, H., Kumar, S.A., Shuai, S., Diaz-Navarro, A., Gutierrez-Fernandez, A., De Antonellis, P., Cavalli, F.M.G., Juraschka, K., Farooq, H., Shibahara, I., et al. (2019). Recurrent noncoding U1 snRNA mutations drive cryptic splicing in SHH medulloblastoma. Nature 574, 707–711.

17. Greene, D., Thys, C., Berry, I.R., Jarvis, J., Ortibus, E., Mumford, A.D., Freson, K., and Turro, E. (2024). Mutations in the U4 snRNA gene RNU4-2 cause one of the most prevalent monogenic neurodevelopmental disorders. Nat Med 30, 2165–2169.

18. Chen, Y., Dawes, R., Kim, H.C., Ljungdahl, A., Stenton, S.L., Walker, S., Lord, J., Lemire, G., Martin-Geary, A.C., Ganesh, V.S., et al. (2024). De novo variants in the RNU4-2 snRNA cause a frequent neurodevelopmental syndrome. Nature 632, 832–840.

19. Pellizzoni, L., Yong, J., and Dreyfuss, G. (2002). Essential role for the SMN complex in the specificity of snRNP assembly. Science 298, 1775–1779.

20. Battle, D.J., Kasim, M., Yong, J., Lotti, F., Lau, C.K., Mouaikel, J., Zhang, Z., Han, K., Wan, L., and Dreyfuss, G. (2006). The SMN complex: an assembly machine for RNPs. Cold Spring Harb Symp Quant Biol 71, 313–320.

21. Cauchi, R.J. (2010). SMN and Gemins: ‘we are family’ … or are we?: insights into the partnership between Gemins and the spinal muscular atrophy disease protein SMN. Bioessays 32, 1077–1089.

22. Zhang, R., So, B.R., Li, P., Yong, J., Glisovic, T., Wan, L., and Dreyfuss, G. (2011). Structure of a Key Intermediate of the SMN Complex Reveals Gemin2’s Crucial Function in snRNP Assembly. Cell 146, 384–395.

23. Sarachan, K.L., Valentine, K.G., Gupta, K., Moorman, V.R., Gledhill, J.M., Bernens, M., Tommos, C., Wand, A.J., and Van Duyne, G.D. (2012). Solution structure of the core SMN-Gemin2 complex. Biochem J 445, 361–370.

24. Grimm, C., Chari, A., Pelz, J.P., Kuper, J., Kisker, C., Diederichs, K., Stark, H., Schindelin, H., and Fischer, U. (2013). Structural basis of assembly chaperone-mediated snRNP formation. Mol Cell 49, 692–703.

25. Veepaschit, J., Viswanathan, A., Bordonné, R., Grimm, C., and Fischer, U. (2021). Identification and structural analysis of the Schizosaccharomyces pombe SMN complex. Nucleic Acids Res 49, 7207–7223.

26. Yong, J., Golembe, T.J., Battle, D.J., Pellizzoni, L., and Dreyfuss, G. (2004). snRNAs contain specific SMN-binding domains that are essential for snRNP assembly. Mol Cell Biol 24, 2747– 2756.

27. Battle, D.J., Lau, C.K., Wan, L., Deng, H., Lotti, F., and Dreyfuss, G. (2006). The Gemin5 protein of the SMN complex identifies snRNAs. Mol Cell 23, 273–279.

28. Lau, C.K., Bachorik, J.L., and Dreyfuss, G. (2009). Gemin5-snRNA interaction reveals an RNA binding function for WD repeat domains. Nat Struct Mol Biol 16, 486–491.

29. Yong, J., Kasim, M., Bachorik, J.L., Wan, L., and Dreyfuss, G. (2010). Gemin5 delivers snRNA precursors to the SMN complex for snRNP biogenesis. Mol Cell 38, 551–562.

30. Baccon, J., Pellizzoni, L., Rappsilber, J., Mann, M., and Dreyfuss, G. (2002). Identification and characterization of Gemin7, a novel component of the survival of motor neuron complex. J Biol Chem 277, 31957–31962.

31. Carissimi, C., Baccon, J., Straccia, M., Chiarella, P., Maiolica, A., Sawyer, A., Rappsilber, J., and Pellizzoni, L. (2005). Unrip is a component of SMN complexes active in snRNP assembly. FEBS Lett 579, 2348–2354.

32. Carissimi, C., Saieva, L., Gabanella, F., and Pellizzoni, L. (2006). Gemin8 is required for the architecture and function of the survival motor neuron complex. J Biol Chem 281, 37009–37016.

33. Otter, S., Grimmler, M., Neuenkirchen, N., Chari, A., Sickmann, A., and Fischer, U. (2007). A comprehensive interaction map of the human survival of motor neuron (SMN) complex. J Biol Chem 282, 5825–5833.

34. Shukla, S., and Parker, R. (2014). Quality control of assembly-defective U1 snRNAs by decapping and 5’-to-3’ exonucleolytic digestion. Proc Natl Acad Sci U S A 111, E3277–86.

35. Prusty, A.B., Meduri, R., Prusty, B.K., Vanselow, J., Schlosser, A., and Fischer, U. (2017). Impaired spliceosomal UsnRNP assembly leads to Sm mRNA down-regulation and Sm protein degradation. J Cell Biol 216, 2391–2407.

36. Lefebvre, S., Bürglen, L., Reboullet, S., Clermont, O., Burlet, P., Viollet, L., Benichou, B., Cruaud, C., Millasseau, P., and Zeviani, M. (1995). Identification and characterization of a spinal muscular atrophy-determining gene. Cell 80, 155–165.

37. Liu, Q., and Dreyfuss, G. (1996). A novel nuclear structure containing the survival of motor neurons protein. EMBO J 15, 3555–3565.

38. Lefebvre, S., Burlet, P., Liu, Q., Bertrandy, S., Clermont, O., Munnich, A., Dreyfuss, G., and Melki, J. (1997). Correlation between severity and SMN protein level in spinal muscular atrophy. Nat Genet 16, 265–269.

39. Wan, L., Battle, D.J., Yong, J., Gubitz, A.K., Kolb, S.J., Wang, J., and Dreyfuss, G. (2005). The survival of motor neurons protein determines the capacity for snRNP assembly: biochemical deficiency in spinal muscular atrophy. Mol Cell Biol 25, 5543–5551.

40. Baserga, S.J., and Steitz, J.A. (1993). The Diverse World of Small Ribonucleoproteins. In The RNA World, R.F. Gesteland, and J.F. Atkins, eds. (Spring Harbor Laboratory Press), pp. 359–381.

41. Mount, S.M., Pettersson, I., Hinterberger, M., Karmas, A., and Steitz, J.A. (1983). The U1 small nuclear RNA-protein complex selectively binds a 5’ splice site in vitro. Cell 33, 509–518.

42. Kaida, D., Berg, M.G., Younis, I., Kasim, M., Singh, L.N., Wan, L., and Dreyfuss, G. (2010). U1 snRNP protects pre-mRNAs from premature cleavage and polyadenylation. Nature 468, 664–668.

43. Berg, M.G., Singh, L.N., Younis, I., Liu, Q., Pinto, A.M., Kaida, D., Zhang, Z., Cho, S., Sherrill-Mix, S., Wan, L., et al. (2012). U1 snRNP determines mRNA length and regulates isoform expression. Cell 150, 53–64.

44. Almada, A.E., Wu, X., Kriz, A.J., Burge, C.B., and Sharp, P.A. (2013). Promoter directionality is controlled by U1 snRNP and polyadenylation signals. Nature 499, 360–363.

45. Preker, P., Nielsen, J., Kammler, S., Lykke-Andersen, S., Christensen, M.S., Mapendano, C.K., Schierup, M.H., and Jensen, T.H. (2008). RNA exosome depletion reveals transcription upstream of active human promoters. Science 322, 1851–1854.

46. Ntini, E., Järvelin, A.I., Bornholdt, J., Chen, Y., Boyd, M., Jørgensen, M., Andersson, R., Hoof, I., Schein, A., Andersen, P.R., et al. (2013). Polyadenylation site-induced decay of upstream transcripts enforces promoter directionality. Nat Struct Mol Biol 20, 923–928.

47. Chiu, A.C., Suzuki, H.I., Wu, X., Mahat, D.B., Kriz, A.J., and Sharp, P.A. (2018). Transcriptional Pause Sites Delineate Stable Nucleosome-Associated Premature Polyadenylation Suppressed by U1 snRNP. Molecular cell 69, 648–663. e7.

48. Oh, J.M., Di, C., Venters, C.C., Guo, J., Arai, C., So, B.R., Pinto, A.M., Zhang, Z., Wan, L., Younis, I., et al. (2017). U1 snRNP telescripting regulates a size-function-stratified human genome. Nat Struct Mol Biol 24, 993–999.

49. Yin, Y., Lu, J.Y., Zhang, X., Shao, W., Xu, Y., Li, P., Hong, Y., Cui, L., Shan, G., Tian, B., et al. (2020). U1 snRNP regulates chromatin retention of noncoding RNAs. Nature 580, 147–150.

50. Nelissen, R.L., Will, C.L., van Venrooij, W.J., and Lührmann, R. (1994). The association of the U1-specific 70K and C proteins with U1 snRNPs is mediated in part by common U snRNP proteins. EMBO J 13, 4113–4125.

51. Pomeranz Krummel, D.A., Oubridge, C., Leung, A.K., Li, J., and Nagai, K. (2009). Crystal structure of human spliceosomal U1 snRNP at 5.5 A resolution. Nature 458, 475–480.

52. Weber, G., Trowitzsch, S., Kastner, B., Lührmann, R., and Wahl, M.C. (2010). Functional organization of the Sm core in the crystal structure of human U1 snRNP. EMBO J 29, 4172–4184.

53. Kondo, Y., Oubridge, C., van Roon, A.M., and Nagai, K. (2015). Crystal structure of human U1 snRNP, a small nuclear ribonucleoprotein particle, reveals the mechanism of 5’ splice site recognition. Elife 4, e04986.

54. So, B.R., Wan, L., Zhang, Z., Li, P., Babiash, E., Duan, J., Younis, I., and Dreyfuss, G. (2016). A U1 snRNP-specific assembly pathway reveals the SMN complex as a versatile hub for RNP exchange. Nat Struct Mol Biol 23, 225–230.

55. Rösel, T.D., Hung, L.-H., Medenbach, J., Donde, K., Starke, S., Benes, V., Rätsch, G., and Bindereif, A. (2011). RNA-Seq analysis in mutant zebrafish reveals role of U1C protein in alternative splicing regulation. EMBO J 30, 1965–1976.

56. Rosel-Hillgartner, T.D., Hung, L.H., Khrameeva, E., Le Querrec, P., Gelfand, M.S., and Bindereif, A. (2013). A novel intra-U1 snRNP cross-regulation mechanism: alternative splicing switch links U1C and U1-70K expression. PLoS Genet 9, e1003856.

57. Muto, Y., Pomeranz Krummel, D., Oubridge, C., Hernandez, H., Robinson, C.V., Neuhaus, D., and Nagai, K. (2004). The structure and biochemical properties of the human spliceosomal protein U1C. J Mol Biol 341, 185–198.

58. White, D.S., Dunyak, B.M., Vaillancourt, F.H., and Hoskins, A.A. (2024). A sequential binding mechanism for 5’ splice site recognition and modulation for the human U1 snRNP. Nat Commun 15, 8776.

59. Hamm, J., van Santen, V.L., Spritz, R.A., and Mattaj, I.W. (1988). Loop I of U1 small nuclear RNA is the only essential RNA sequence for binding of specific U1 small nuclear ribonucleoprotein particle proteins. Mol Cell Biol 8, 4787–4791.

60. Cheng, D., Côté, J., Shaaban, S., and Bedford, M.T. (2007). The arginine methyltransferase CARM1 regulates the coupling of transcription and mRNA processing. Mol Cell 25, 71–83.

61. Friesen, W.J., Massenet, S., Paushkin, S., Wyce, A., and Dreyfuss, G. (2001). SMN, the product of the spinal muscular atrophy gene, binds preferentially to dimethylarginine-containing protein targets. Mol Cell 7, 1111–1117.

62. Selenko, P., Sprangers, R., Stier, G., Bühler, D., Fischer, U., and Sattler, M. (2001). SMN tudor domain structure and its interaction with the Sm proteins. Nat Struct Biol 8, 27–31.

63. Brahms, H., Meheus, L., de Brabandere, V., Fischer, U., and Lührmann, R. (2001). Symmetrical dimethylation of arginine residues in spliceosomal Sm protein B/B’ and the Sm-like protein LSm4, and their interaction with the SMN protein. RNA 7, 1531–1542.

64. Lerner, M.R., Boyle, J.A., Mount, S.M., Wolin, S.L., and Steitz, J.A. (1980). Are snRNPs involved in splicing. Nature 283, 220–224.

65. Zhuang, Y., and Weiner, A.M. (1986). A compensatory base change in U1 snRNA suppresses a 5’ splice site mutation. Cell 46, 827–835.

66. Sharma, S., Maris, C., Allain, F.H., and Black, D.L. (2011). U1 snRNA directly interacts with polypyrimidine tract-binding protein during splicing repression. Mol Cell 41, 579–588.

67. Duckett, D.R., Murchie, A.I., and Lilley, D.M. (1995). The global folding of four-way helical junctions in RNA, including that in U1 snRNA. Cell 83, 1027–1036.

68. Hamm, J., Dathan, N.A., Scherly, D., and Mattaj, I.W. (1990). Multiple domains of U1 snRNA, including U1 specific protein binding sites, are required for splicing. The EMBO journal 9, 1237.

69. Grimmler, M., Bauer, L., Nousiainen, M., Körner, R., Meister, G., and Fischer, U. (2005). Phosphorylation regulates the activity of the SMN complex during assembly of spliceosomal U snRNPs. EMBO Rep 6, 70–76.

70. Petri, S., Grimmler, M., Over, S., Fischer, U., and Gruss, O.J. (2007). Dephosphorylation of survival motor neurons (SMN) by PPM1G/PP2Cgamma governs Cajal body localization and stability of the SMN complex. J Cell Biol 179, 451–465.

71. Schilling, M., Prusty, A.B., Boysen, B., Oppermann, F.S., Riedel, Y.L., Husedzinovic, A., Rasouli, H., König, A., Ramanathan, P., Reymann, J., et al. (2021). TOR signaling regulates liquid phase separation of the SMN complex governing snRNP biogenesis. Cell Rep 35, 109277.

72. Raimer, A.C., Gray, K.M., and Matera, A.G. (2016). SMN - A chaperone for nuclear RNP social occasions. RNA Biol 14, 701–711.

73. Šimčíková, D., Gelles-Watnick, S., and Neugebauer, K.M. (2023). Tudor-dimethylarginine interactions: the condensed version. Trends Biochem Sci 48, 689–698.

74. Lanfranco, M., Vassallo, N., and Cauchi, R.J. (2017). Spinal Muscular Atrophy: From Defective Chaperoning of snRNP Assembly to Neuromuscular Dysfunction. Front Mol Biosci 4, 41.

75. Kour, S., Rajan, D.S., Fortuna, T.R., Anderson, E.N., Ward, C., Lee, Y., Lee, S., Shin, Y.B., Chae, J.H., Choi, M., et al. (2021). Loss of function mutations in GEMIN5 cause a neurodevelopmental disorder. Nat Commun 12, 2558.

76. Saida, K., Tamaoki, J., Sasaki, M., Haniffa, M., Koshimizu, E., Sengoku, T., Maeda, H., Kikuchi, M., Yokoyama, H., Sakamoto, M., et al. (2021). Pathogenic variants in the survival of motor neurons complex gene GEMIN5 cause cerebellar atrophy. Clin Genet 100, 722–730.

77. Earnest-Noble, L.B., Hsu, D., Chen, S., Asgharian, H., Nandan, M., Passarelli, M.C., Goodarzi, H., and Tavazoie, S.F. (2022). Two isoleucyl tRNAs that decode synonymous codons divergently regulate breast cancer metastatic growth by controlling translation of proliferation-regulating genes. Nat Cancer 3, 1484–1497.

78. Goraczniak, R., Behlke, M.A., and Gunderson, S.I. (2009). Gene silencing by synthetic U1 adaptors. Nat Biotechnol 27, 257–263.

79. Fernandez Alanis, E., Pinotti, M., Dal Mas, A., Balestra, D., Cavallari, N., Rogalska, M.E., Bernardi, F., and Pagani, F. (2012). An exon-specific U1 small nuclear RNA (snRNA) strategy to correct splicing defects. Hum Mol Genet 21, 2389–2398.

80. Gonçalves, M., Santos, J.I., Coutinho, M.F., Matos, L., and Alves, S. (2023). Development of Engineered-U1 snRNA Therapies: Current Status. Int J Mol Sci 24, 14617.

81. Lee, S.H., Singh, I., Tisdale, S., Abdel-Wahab, O., Leslie, C.S., and Mayr, C. (2018). Widespread intronic polyadenylation inactivates tumour suppressor genes in leukaemia. Nature 561, 127–131.

82. Hsu, T.Y., Simon, L.M., Neill, N.J., Marcotte, R., Sayad, A., Bland, C.S., Echeverria, G.V., Sun, T., Kurley, S.J., Tyagi, S., et al. (2015). The spliceosome is a therapeutic vulnerability in MYC-driven cancer. Nature 525, 384–388.

83. Love, S.L., Emerson, J.D., Koide, K., and Hoskins, A.A. (2023). Pre-mRNA splicing-associated diseases and therapies. RNA Biol 20, 525–538.

84. Ji, P., Diederichs, S., Wang, W., Böing, S., Metzger, R., Schneider, P.M., Tidow, N., Brandt, B., Buerger, H., Bulk, E., et al. (2003). MALAT-1, a novel noncoding RNA, and thymosin beta4 predict metastasis and survival in early-stage non-small cell lung cancer. Oncogene 22, 8031–8041.

85. Esteller, M. (2011). Non-coding RNAs in human disease. Nat Rev Genet 12, 861–874.

86. Goodarzi, H., Liu, X., Nguyen, H.C., Zhang, S., Fish, L., and Tavazoie, S.F. (2015). Endogenous tRNA-Derived Fragments Suppress Breast Cancer Progression via YBX1 Displacement. Cell 161, 790–802.

87. Oh, J.M., Venters, C.C., Di, C., Pinto, A.M., Wan, L., Younis, I., Cai, Z., Arai, C., So, B.R., Duan, J., et al. (2020). U1 snRNP regulates cancer cell migration and invasion in vitro. Nat Commun 11, 1.

88. Munding, E.M., Shiue, L., Katzman, S., Donohue, J.P., and Ares, M. (2013). Competition between pre-mRNAs for the splicing machinery drives global regulation of splicing. Mol Cell 51, 338–348.

89. Kumar, P., Joy, J., Pandey, A., and Gupta, D. (2017). PRmePRed: A protein arginine methylation prediction tool. PLoS One 12, e0183318.

90. Wan, L., Ottinger, E., Cho, S., and Dreyfuss, G. (2008). Inactivation of the SMN complex by oxidative stress. Mol Cell 31, 244–254.

